# A nuclear TRiC/CCT chaperonin assembles meiotic HORMAD proteins into chromosome axes competent for crossing over

**DOI:** 10.1101/2025.03.03.641182

**Authors:** Monique Zetka, Amirhossein Pezeshki, Steven J.M. Jones, Nicola Silva

## Abstract

The meiotic chromosome axis organizes chromatin and sets the stage for homolog pairing and recombination. <u>M</u>eiotic <u>HORMA</u> <u>d</u>omain proteins (mHORMADs) are conserved axis components that conformationally transform during target binding. In *C. elegans,* four functionally distinct mHORMADs directly interact, but how binding between them is restricted to axis assembly is unknown. Using a mutation in the mHORMADs that delays axis assembly, we isolated a suppressor mutation in a TRiC/CCT chaperonin subunit that restored mHORMAD localization. CCT-4 associates with meiotic chromatin and forms *in vivo* complexes with mHORMADs, while germline disruption of TRiC results in axis defects, indicating a nuclear function for TRiC alongside meiotic chromosomes. We propose that chromosome-tethered TRiC folds mHORMADs into a conformationally active local population required for axis morphogenesis. More broadly, our results support the model that spatially-restricted folding by TRiC/CCT is a mechanism of controlling the assembly of multimeric complexes that function in tightly co-ordinated events.

## Introduction

A central feature of meiosis is the formation of crossovers that direct the accurate segregation of chromosomes at the first division. <u>C</u>ross<u>o</u>ver (CO) formation depends on earlier events that pair homologous chromosomes together and initiate recombination through the formation of meiotic <u>d</u>ouble-<u>s</u>trand <u>b</u>reaks (DSBs) that are repaired by <u>h</u>omologous recombination (HR) ^1^. How homology is assessed between chromosomes is an enduring mystery; however, once established, homologs juxtapose along their lengths, and their paired configuration is stabilized by the formation of the <u>s</u>ynaptonemal <u>c</u>omplex (SC). The SC is an ordered assembly of <u>c</u>entral region (CR) proteins that polymerize between the axes of two chromosomes where they promote interhomolog recombination and CO formation. While the resulting tripartite structure is widely conserved at the ultrastructural level ^2^, the individual CR components that make up the central region in different organisms share little sequence similarity ^3,4^.

Previous studies in a variety of organisms have shown that synapsis *per se* does not distinguish homology since SC forms between nonhomologous DNA sequences ^5^. CR components have an innate proclivity for self-assembly and can readily polymerize in situations where the meiotic axes cannot support synapsis, instead forming ordered nuclear polycomplexes that ultrastructurally resemble the SC without the involvement of chromosomes ^6^. Characterization of wild-type SC dynamics have observed that SC assembly is highly processive and rapid once initiated ^7,8^; consequently, nucleation of SC components at chromosome axes must be tightly coordinated with homolog pairing to reduce the risk of ectopic synapsis between nonhomologous sequences. In most experimental meiotic model systems, this problem is solved by SC nucleation at sites of homologous recombination between paired homologs; alternatively, specific chromosomal sites are used for both pairing and initiation of SC assembly, independent of DSB formation and the HR pathway ^9^. The DSB-independent pathway is most clearly illustrated by *C. elegans,* where each chromosome contains a <u>p</u>airing <u>c</u>enter (PC) that recruits one member of a family of DNA-binding proteins required for pairing (*zim-1, 2, 3* and *him-8* ^10,11^), and is also the preferred site of synapsis initiation ^7,12,13^. PC regions are tethered to the nuclear envelope where they connect the chromosome to cytoplasmic forces that drive pairing ^13–17^, promote SC assembly, and eliminate interactions between nonhomologs that could lead to synapsis between them ^7,13,14,17–19^. The assembly of the PC, pairing, synapsis, and recombination are all dependent on the earlier formation of the meiotic chromosome axis, indicating that axis proteins are required for configuring the chromosome for meiotic processes ^20–24^.

The composition of the meiotic axes of yeasts, animals, and plants is dominated by cohesin complexes and member(s) of a family of conserved meiosis-specific proteins containing a HORMA domain (mHORMAD proteins) ^25–27^. The HORMAD is a structural feature first identified in <u>Ho</u>p1p, <u>Re</u>v7p and <u>M</u>ad2p during systemic analysis of budding yeast DNA repair proteins ^28^. Early structural and biochemical studies of the mitotic checkpoint protein Mad2 revealed that the HORMA domain was structurally metamorphic and could reversibly interconvert between an open/active and closed/inactive conformation, a dramatic structural transition in which the HORMAD core encircles a “safety belt” in the C-terminal region around a peptide sequence on a binding partner ^29^. Unlike Mad2 however, no stable open conformation has been observed for any mHORMAD protein ^30,31^ and they instead adopt a partially unfolded form in which the safety belt is “unbuckled” from the intact HORMA domain core ^32^. The meiotic HORMAD proteins have essential functions at chromosome axes in DSB formation, homolog pairing, SC formation, and CO formation ^27^, and are represented by Hop1 in *S. cerevisiae* ^33–36^, HORMAD1 and HORMAD2 in mice ^37–40^, and ASY1 in plants ^41^. *C. elegans* has four mHORMAD proteins (HIM-3, HTP-1, HTP-2, HTP-3) that interact to assemble into chromosome axes ^23, 42, 43^. HTP-3 binds first, possibly to a sequence within the meiotic cohesin complex ^44^ and then recruits HIM-3 and HTP-1/2 ^23,45,46^ which use their HORMA domains to bind to multiple short sequences located in the extended HTP-3 C-terminal tail and switch to closed forms ^42^. Despite their direct physical interactions, the mHORMADs have distinct functions in meiotic processes. HTP-3 is required for meiotic DSB formation and <u>s</u>ister <u>c</u>hromatid <u>c</u>ohesion (SCC ^46,47^), while HIM-3 is required for homolog pairing, SC formation, and the bias to using the homolog as a DSB repair template ^48,49^. HTP-1 and HTP-2 (referred to as HTP-1/2) are highly homologous (>80% a.a. identity,) but show little functional redundancy; while *htp-2* mutants have no obvious phenotype, *htp-1* mutants show severe defects in synapsis and CO formation that are exacerbated by loss of *htp-2*. ^21,22^ The abundant nonhomologous synapsis observed in *htp-1* mutants is accompanied by severe pairing defects, indicating that HTP-1 plays a regulatory role that restricts SC assembly to paired PCs ^15,21,22^. Recent work has identified a conserved extended loop in the safety belt of the mHORMAD that could expand the range of possible conformations and create structural signatures for protein interactions that could explanation their functional specificity ^50^.

Free mHORMAD proteins that are not bound to their binding partner are thought to exist primarily in an inactive self-closed state by interacting with their own C-terminal domain, and consequently require remodelling to convert into the active unbuckled conformation ^31, 43, 50^. In yeasts, mammals, and plants, mHORMAD localization dynamics depend on Pch2/TRIP, a conserved checkpoint AAA-ATPase that can unbuckle closed conformations of the mHORMADs to produce binding-competent-pools for loading onto chromosomes, and to remove or redistribute axis-bound mHORMADs in response to progression of meiotic cell-cycle events.^29,51,52^ Consistent with functions in regulating mHORMAD chromosome association, Pch2 orthologs in budding yeast^53^, plants^54,55^, and mice ^56^ localize to chromosomes upon meiotic entry and then colocalize with the SC into pachytene stages. *C. elegans* PCH-2 similarly appears first as foci on meiotic chromosomes, followed by localization to the SC; however, the localization of the mHORMADs is PCH-2-independent^57^, indicating that their remodelling into a binding-competent form during axis assembly is mediated by an unknown pathway.

Molecular chaperones are required to assist in the folding of cellular proteins by recognizing and binding newly synthesized polypeptides to prevent premature folding and aggregation. The eukaryotic <u>T</u>ailless complex peptide 1 <u>Ri</u>ng <u>C</u>omplex (TRiC) or <u>c</u>haperonin-<u>c</u>ontaining <u>T</u>-complex protein 1 (CCT) complex is a large type II chaperonin that uses an ATP hydrolysis-driven cycle to fold proteins in a central cavity formed by two stacked rings, each containing 8 highly related CCT subunits.^58^ TRiC substrates are a structurally diverse population with complex topologies that cannot be folded by the simpler chaperone system, and as a general rule contain extended hydrophobic stretches (particularly β-sheets), are aggregation prone, and participate in multiprotein complexes ^59, 60^. In addition to its obligate cytosolic substrates actin and tubulin, TRiC is also required for the folding of nuclear proteins that function in cell-cycle regulation, transcription, chromatin modification, and genome stability.^59,61^ Although this complex has historically been considered to function exclusively as a cytoplasmic factor, nuclear TRiC has been identified using proteomic approaches ^62^ and studies across a broad range of systems have reported nuclear localization of TRiC subunits ^63–68^, that suggested the existence of TRiC-mediated functions within the nucleus. Moreover, CCT subunits localize to germline nuclei and with the chromatin of rat spermatocytes,^69^ consistent with functions in the germline compartment that extend to chromosome biology. The existence of active protein folding machinery working alongside the genome is not without precedent. The major molecular chaperones (Hsp90, Hsp70, and Hsp60) are known to participate in the stabilization of nuclear proteins, and in the assembly and disassembly of complexes functioning in replication, transcription and other DNA-related pathways^70^. A nuclear function for TRiC in modulating RNAPII activity has been reported in budding yeast, indicating that chaperonins may similarly be required to locally regulate assembly and disassembly of nuclear multisubunit complexes^71^.

In this study, we report an essential nuclear function for TRiC in meiotic axis morphogenesis. *cct-4*, the δ subunit of the nematode TRiC/CCT complex, interacts with mHORMAD proteins *in vivo* and is required for their axis localization. These results demonstrate an essential role for the chaperonin in regulating meiotic processes leading to CO formation. Based on these data and other evidence, we propose that nuclear TRiC is tethered to meiotic chromosomes where it is required to convert self-closed mHORMAD proteins into an active unbuckled conformation that can bind to interacting proteins during axis assembly. Our results suggest that the complex protein interaction network that is the basis of meiotic axis assembly is regulated at the level of local mHORMAD protein folding and release in an active conformation, consistent with a model of just-in-time protein folding by TRiC^72^ as a mechanism of controlling tightly co-ordinated events in meiotic prophase.

## Results

To identify the pathways regulating the assembly of chromosome axes, we made use of a unique missense allele of *him-3* that results in a S35F substitution at a widely conserved serine within the HORMA domain (Fig. 1A,B). *him-3(vv6)* is expressed at normal levels and the protein localizes to axes, but mutants exhibit severe pairing defects that are accompanied by extensive synapsis between nonhomologous chromosomes (^49^; this study). Upon closer examination, we discovered that *him-3(vv6)* mutants exhibited delayed loading of HIM-3 (described in detail in the next section) and used the allele in a suppressor screen to identify genetic pathways that function in axis assembly. The high embryonic lethality that accompanied the loss of crossing over in *him-3(vv6)* mutants was exploited to perform an EMS-based screen to isolate mutants that increased the brood size. We recovered 4 dominant extragenic suppressors (*vv39, vv41, vv50, vv52*) that increased the number of progeny produced by *him-3(vv6)* mutants 3-6 fold, and focused on *vv39*, the strongest suppressor. SNP mapping^73^ and sequencing revealed a single C to T mutation that mapped to the third exon of the *cct-4* gene. To confirm that *cct-4(vv39)* corresponded to the suppressor mutation, a single copy germline expression construct of *cct-4(vv39)* was integrated into chromosome II using MosSCI^74^ and tested for its ability to dominantly suppress *him-3(vv6)* phenotypes. While *him-3(vv6)* mutants had an average progeny number of 23<u>^+^</u>11, *vvIs17[pie-1p::cct-4(vv39)::unc-54utr]; him-3(vv6)* homozygotes produced 131±34, a value not significantly different from the average progeny number observed for the original *him-3(vv6); cct-4(vv39)* suppressed line (103±42). Examination of the diakinesis nuclei of *him-3(vv6); vvIs17* mutant germlines revealed fewer DAPI-stained bodies in comparison to *him-3(vv6)* mutants that were not significantly different from *him-3(vv6); cct-4(vv39)* mutant germlines, indicating an increase in the number of bivalents (Fig. 1C). These results indicate that both endogenous and transgenic *cct-4(vv39)* expression in the germline can partially suppress the defects in chiasma formation and embryonic lethality associated with *him-3(vv6)* mutants, and demonstrate that *cct-4(vv39)* is the suppressor mutation. *cct-4* encodes the ortholog of human CCT4, the δ subunit of the conserved eukaryotic type II chaperonin; *vv39* corresponds to a missense mutation that results in a P382S substitution in the predicted protein coding sequence. P382 is conserved in CCT4 subunits across species, but is not found in the other nematode TRiC subunits, indicating that it is a conserved feature specific to CCT-4 (Fig. S1). The TRiC/CCT complex is formed by two stacked rings, each composed of eight paralogous CCT subunits^75^. Each subunit has a tripartite organization that contributes to the overall function of the complex: the equatorial domain contains the site of ATP binding, an intermediate domain acts as a hinge during opening and closing of the complex, and an apical domain is generally considered responsible for substrate recognition and binding^58^. In the case of *cct-4(vv39),* the affected proline (P382S) sits at the junction of the apical and intermediate domains (Fig. 1B), suggesting that *vv39* will affect how the chaperonin interacts with polypeptides, rather than regulation by ATP or the overall double ring structure.

**Figure 1.**
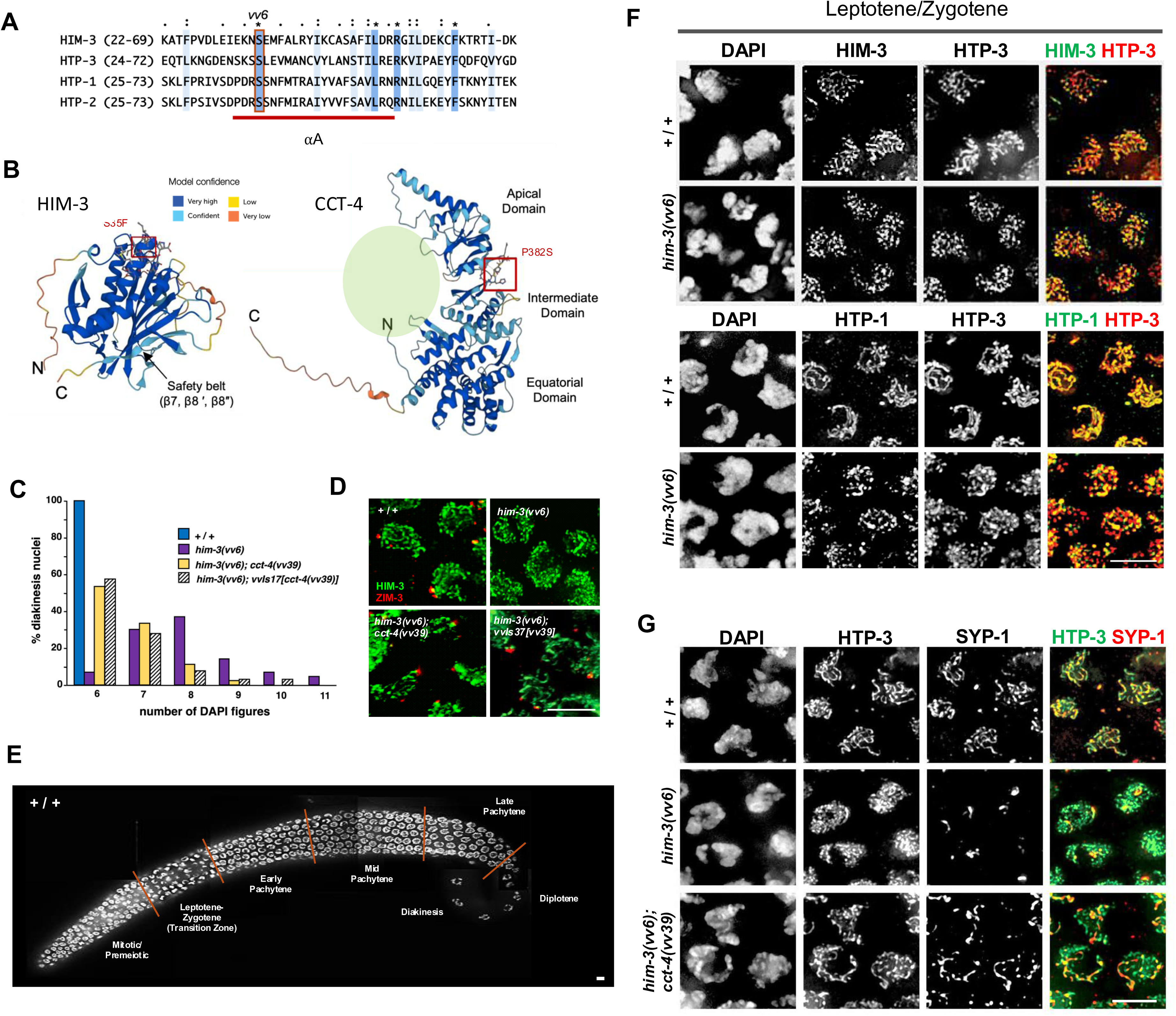
Delayed axis assembly in *him-3(vv6)* mutants is rescued by *cct-4(vv39)*. **A)** M-Coffee multiple sequence alignment of the *C. elegans* meiotic HORMAD proteins showing the N-terminal portion of the HORMA domain and the position of the *vv6* mutation. **B)** AlphaFold structural prediction^117^ of wild-type HIM-3 and CCT-4 showing the position of S35 and P382, the residues affected by *vv6* and *vv39*, respectively. Green sphere represents TRiC cavity. **C)** Histogram showing the percentage of diakinesis nuclei with the indicated number of DAPI-stained bodies. Wild-types consistently exhibit six DAPI figures representing the six chromosome pairs joined by chiasmata, while *him-3(vv6)* mutants rarely exhibit these levels (p<0.0001)*. cct-4(vv39)* and germline expression of *vvIs17*-encoded CCT-4^P382S^ increases the frequency of appearance of bivalents in *him-3(vv6)* mutant nuclei (p<0.0001 for 6 DAPI bodies for both genotypes); differences between *him-3(vv6); cct-4(vv39)* and *him-3(vv6); vvIs17* are not statistically significant in all cases. **D)** Early transition zone nuclei of the indicated genotypes stained for DNA, HTP-3 and the pairing center binding protein ZIM-3 (localizes to chromosomes I and IV^11^). **E)** DAPI-stained wild-type germlines showing nuclei at different stages of meiotic prophase; the asymmetrically positioned chromatin cluster typical of leptotene-zygotene stages occurs in the nuclei in the transition zone (TZ). **F)** Immunofluorescence micrographs showing localization of DNA and the axis components HIM-3, HTP-3 and HTP-1 in early transition zone nuclei of the indicated genotypes. **G)** Early transition zone nuclei in wild-type, *him-3(vv6)* and *him-3(vv6); cct-4(vv39)* mutant germlines stained for DNA, HTP-3, and the SC component SYP-1. Scale bars, 5µm.

### *him-3(vv6)* mutants show delayed axis morphogenesis that is rescued by *cct-4(vv39)*

To understand how a mutation in a subunit of the TRiC chaperonin could suppress meiotic defects associated with an axis component, we re-examined in detail early pairing stages in *him-3(vv6)* mutants using markers not available at the time of its initial characterization. The germline of *C. elegans* is organized along a temporal-spatial axis in which meiotic nuclei passing through the stages of meiotic prophase can be cytologically distinguished (Fig. 1E). The transition <u>z</u>one (TZ) of the germline is populated by leptotene/zygotene stage nuclei in which chromosomes undergoing pairing are clustered to one side, a polarized arrangement that is dispersed with successful homolog pairing, and stabilized by synapsis during pachytene^76,77^. In the earliest polarized nuceli of wild-type transition zones (denoting meiotic entry), HIM-3 already appeared as continuous lines of variable length and intensity that overlapped with the HTP-3-marked chromosomes (Fig. 1F). In nuclei at the same stage in *him-3(vv6)* mutants, however, HIM-3^S35F^ showed a disrupted localization pattern indicative of a defect in timely chromosome recruitment, often appearing partially diffuse and accompanied by foci of heterogenous sizes and intensities that occasionally formed small segments and grossly overlapped with HTP-3 and chromatin (Fig. 1D,F). These HIM-3^S35F^ localization dynamics resembled those reported for HTP-3 mutants lacking the full complement of HIM-3 binding sites^43^, consistent with the interpretation that *him-3vv6)* mutants are impaired in the binding of HIM-3 to HTP-3, thereby delaying the appearance of HIM-3-marked axes.

HTP-3 is associated with the chromatin of pre-meiotic nuclei, but upon meiotic prophase entry its localization rapidly transforms into robust linear stretches that colocalize with HIM-3 and HTP-1/2^45,47^. HTP-3 appeared at assembling meiotic chromosome axes on time in *him-3(vv6)* mutants; however, in polarized nuclei of the early TZ it first appeared as puncta and thin discontinuous stretches that contrasted with the long HTP-3-marked linear elements observed in wild-type nuclei at the same stage (Fig.1 F,G). Similarly, HTP-1/2 were recruited to chromosome axes in *him-3(vv6)* mutants (Fig. 1F), but in early TZ nuclei also showed an interrupted pattern of localization that eventually became continuous as prophase progressed. The defects in early HTP-3 and HTP-1/2 dynamics observed in *him-3(vv6)* mutants were unexpected given that the other mHORMAD proteins are not dependent on HIM-3 for their localization^43,47^, and instead suggests a role for HIM-3 in the polymerization of axis components into linear structures. This is consistent with the results of an analysis of the four HTP-3 binding sequences (“closure motifs”) that recruit HIM-3; the binding of HIM-3 to a single closure motif could support well-defined axes with uninterrupted stretches of HTP-3 and HTP-1/2 localization (but not synapsis), but complete loss of HIM-3 binding resulted in punctate and disordered HTP-3 and HTP-2 chromosome localization.^42,43^ Consequently, the disorganized axes observed in *him-3(vv6)* mutants are likely a reflection of impaired binding of HIM-3 to its closure motifs on HTP-3, which in turn visibly affects axis polymerization. To confirm the delayed axis association of HIM-3 in *him-3(vv6)* mutants, we took advantage of the fact that SC assembly is highly sensitive to HIM-3 levels at the axis^43^ and used the SC component SYP-1 as an independent marker of HIM-3 dynamics. Consistent with the impaired early localization of HIM-3, SYP-1 also showed a delay in chromosome localization in *him-3(vv6)* mutants; in early TZ nuclei, SYP-1 appeared as 1-2 chromosome-associated aggregates or short chromosome-associated stretches, in comparison to the long axis associated stretches of SYP-1 that were observed in wild-type nuclei at the same stage (Fig. 1G). These results indicated that the mutant *him-3(vv6)* protein is delayed in its association with chromosomes in early prophase, leading to defects in axis assembly that are later evident as defects in synapsis and CO formation (this study; ^49^).

We next examined the effect of *cct-4(vv39)* on the early prophase defects of *him-3(vv6)* mutants and observed a visible rescue of the delayed axis morphogenesis defect; in early TZ nuclei, HTP-3-marked chromosome axes were longer and more continuous in appearance in comparison to *him-3(vv6)* mutants, and long SYP-1 stretches appeared at the axes (Fig. 1G). Since the delayed HIM-3 loading in *him-3(vv6)* mutants is accompanied by severe pairing defects and nonhomologous synapsis (^49^; this study), we conclude that *vv39*-mediated suppression of *him-3(vv6)* defects occurred at the level of improved *him-3* localization that in turn improved timely axis assembly and the downstream events required for CO formation.

### *cct-4(vv39)* restores autosomal PC protein loading and establishment of connections to cytoskeletal forces in *him-3(vv6) mutants*

HIM-3 is required for pairing and synapsis of all chromosomes and localization of PC binding proteins to the autosomes (ZIM-1,2,3); the X chromosome recruits its PC binding protein (HIM-8) independently of HIM-3, but does not pair or synapse.^10,49^ To investigate the origin of the autosomal pairing defect in *him-3(vv6)* mutants, the assembly of PCs was assessed by examining the localization of ZIM-3, which localizes to the PC ends of chromosomes I and IV (Fig. 1D;^11^). In wild-type germlines, ZIM-3 localized to the PCs at the nuclear periphery in TZ nuclei and two bright foci were often observed, corresponding to the paired autosomes (Fig 1D). In *him-3(vv6)* mutants, ZIM-3 was not detectable at any stage, suggesting that insufficient levels of HIM-3 were associated with the axes during PC binding protein recruitment stages. In the presence of *cct-4(vv39)* and *vvIs17*, however, ZIM-3 reappeared as foci at the nuclear periphery of polarized TZ nuclei in the suppressed germlines (Fig. 1D). Since ZIM-3 recruitment to PCs is defective in *him-3(vv6)* mutants, these results suggest that configuring the PC for ZIM binding is disrupted by the earlier delay in axis morphogenesis and can be restored by improved HIM-3^S35F^ association with HTP-3 in the presence of CCT-4^P382S^.

During chromosome pairing stages, cytoplasmic forces are transmitted through an intact NE to the ends of meiotic chromosomes to generate motions that facilitate pairing and discourage nonhomologous synapsis.^78^ In *C. elegans*, only the PC end of the chromosome interacts with SUN-1/Matefin, the inner NE membrane protein that couples with ZYG-12 to bridge the NE and connect to dynein-driven forces; PC binding proteins at the PCs recruit Polo-like kinase 2 (PLK-2), which is required for local aggregation of SUN-1/ZYG-12 and the concentration of forces at the chromosome end to generate movement.^3–15,79,80^. Given the loss of ZIM-3 localization in *him-3(vv6*) mutants, we used PLK-2 as a proxy for PC binding protein localization to investigate if the pairing defects in *him-3(vv6)* mutants could be explained by a global failure in autosomal PC binding protein localization and loss of the PLK-2 activity required for chromosome movement (Fig. 2A). In *him-3(vv6)* mutants, PLK-2 localization was reduced to a bright focus that correlated to the X chromosome PC, indicating a failure in autosomal PC binding protein localization. In *him-3(vv6); cct-4(vv39)* suppressed germlines however, numerous larger aggregates of PLK-2 formed at the nuclear periphery (representing clusters of variable numbers of chromosome ends), resembling the 4-6 aggregates that are observed in wild types.^79,80^ To confirm that the PLK-2 aggregates were functional, we monitored the presence of SUN-1 phosphorylation at S12 (Fig. 2B), a PLK-2-dependent post-translational modification specifically associated with the SUN-1 population aggregated around PCs.^14^ In wild types, SUN-1^S12-Pi^ appears as 1-3 variably sized aggregates in transition zone nuclei^14,79,80^, however, in *him-3(vv6)* mutant germlines only one SUN-1^S12-Pi^ aggregate was observed, and this colocalized with HIM-8 (Fig. 2B). In contrast, 1-3 patches of SUN-1^S12-Pi^ were observed in *him-3(vv6); cct-4(vv39*) suppressed germlines, and the TZ nuclei appeared more tightly clustered than in *him-3(vv6)* mutants, collectively consistent with improved chromosome movement at pairing stages. Similarly, localization ZYG-12 revealed a single ZYG-12 patch per nucleus in *him-3(vv6)*, while in *him-3(vv6); cct-4(vv39)* suppressed germlines, multiple ZYG-12 patches were formed (Fig. 2C). In summary, we conclude that the links between autosomal PCs and the cytoskeletal forces required for homolog pairing is defective in *him-3(vv6)* mutants, and is partially restored in the presence of *cct-4(vv39)*. Based on these data, however, we cannot distinguish if *cct-4(vv39)* restores PC binding protein localization to all autosomes and the partial suppression is a consequence of a downstream defect, or if each autosome is variably affected by *vv39*-mediated suppression of the axis morphogenesis defects; those that can localize the PC proteins establish SUN-1 linkages while those that fail do not.

**Figure 2.**
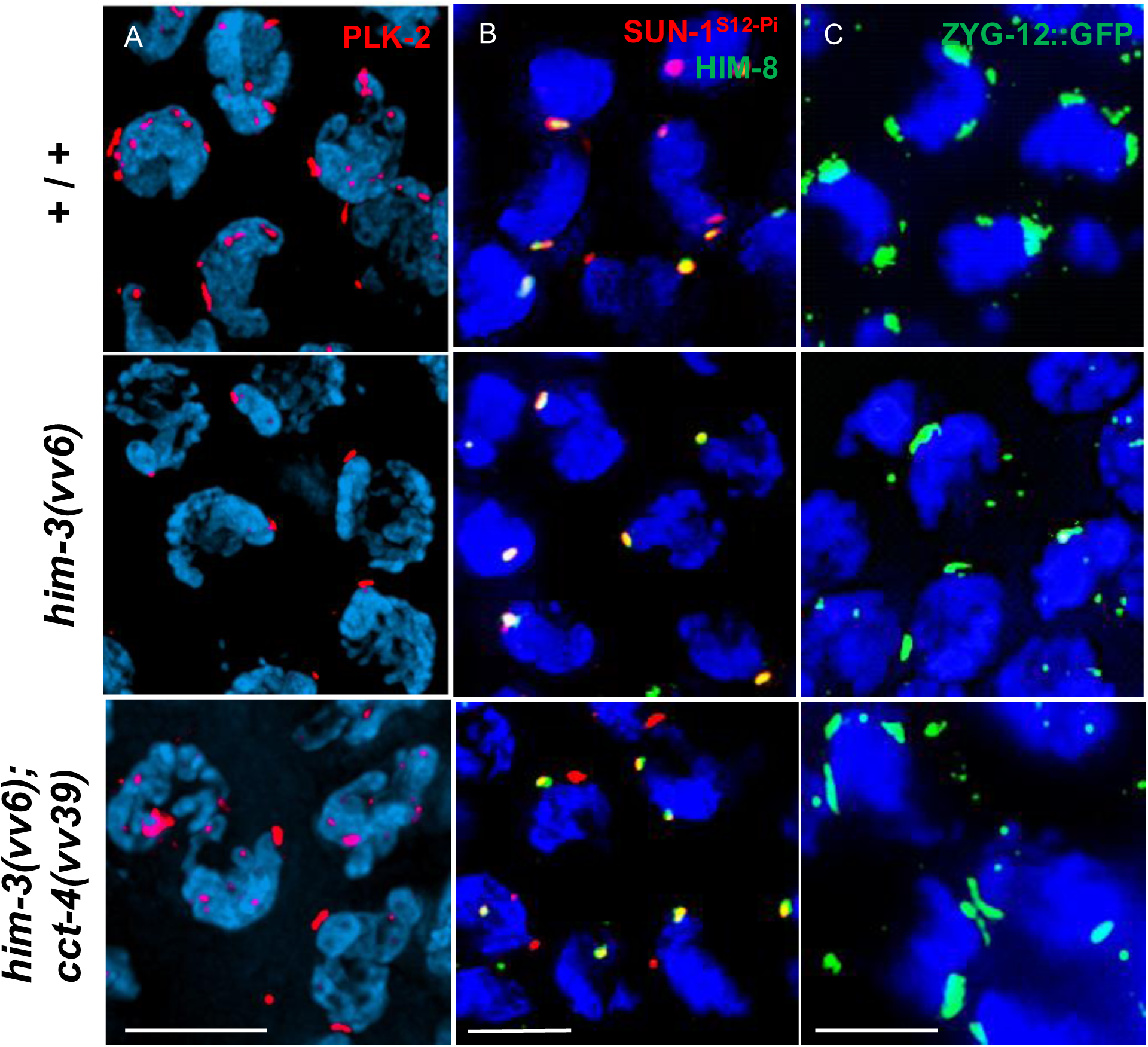
Autosomal linkages to cytoskeletal forces missing in *him-3(vv6)* mutants reappear in the presence of *cct-4(vv39)*. Immunofluorescence micrographs of DAPI-stained transition zone nuclei from germlines of the indicated genotypes showing localization of **A)** the polo-like kinase PLK-2, **B)** the X chromosome PC binding protein HIM-8 and the population of SUN-1 phosphorylated at S12, and **C)** the outer NE component ZYG-12 (detected by α-GFP). Scale bars, 5µm.

### *cct-4(vv39)* partially restores chromosome pairing and its coordination with synapsis in *him-3(vv6)* mutants

Since *cct-4(vv39)* suppressed both the timely axis morphogenesis and ZIM-3 localization defects of *him-3(vv6)*, we next investigated if these events were accompanied by an improvement in chromosome pairing using fluorescence *in <u>s</u>itu* <u>h</u>ybridization (FISH) to assay pairing of the 5S rDNA locus in a time-course analysis (Fig. 3A). In wild-type germlines, the percentage of nuclei with paired FISH signals jumped to 55% upon entry into leptotene/zygotene (p<0.0001), and reached 99% by late pachytene (p<0.0001), indicative of homologous pairing stabilized by full synapsis^81^. Consistent with the findings of a previous study, *him-3(vv6)* mutant showed severe pairing defects from the earliest stages of meiotic prophase^49^. The level of pairing at the beginning of leptotene/zygotene in *him-3(vv6)* mutant germlines was not significantly different in comparison to the premeiotic region (9%, p=0.25); but reached 18% by the end of the extended transition zone (p*=*0.02). In suppressed *him-3(vv6); cct-4(vv39)* germlines, pairing at the beginning of leptotene/zygotene remained at *him-3(vv6)* mutant levels (p=0.81), but steadily rose to 47% by late pachytene, 2.6-fold higher than the levels observed in *him-3(vv6)* mutant nuclei at the same stage (p<0.0001). These results indicated that the early pairing defect of *him-3(vv6)* mutants persisted in *him-3(vv6); cct-4(vv39)* germlines, but significantly improved as prophase progressed.

**Figure 3.**
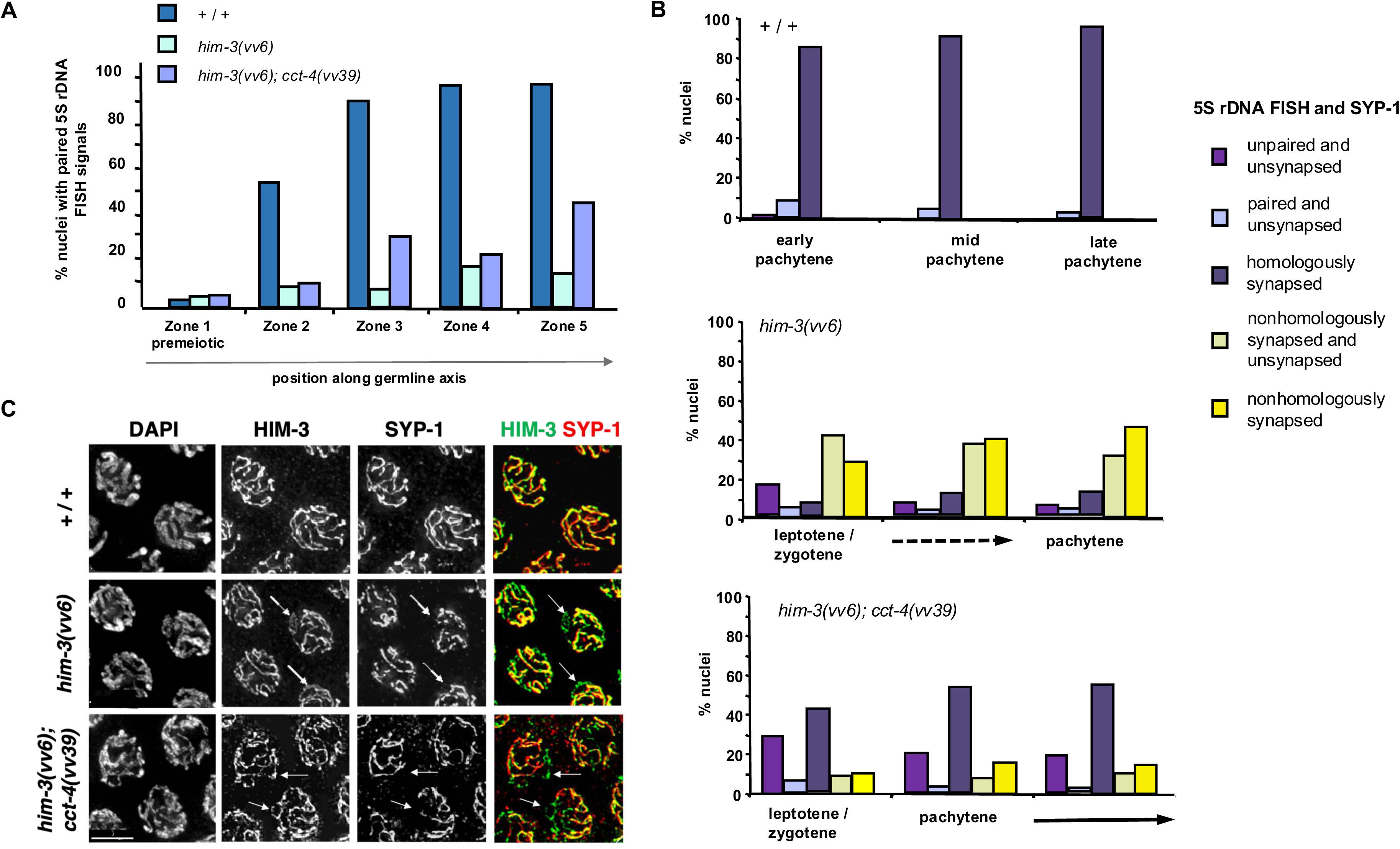
*cct-4(vv39)* partially suppresses the homologue pairing defects of *him-3(vv6)* mutants. **A)** Histograms showing a time-course analysis of pairing levels in wild types, *him-3(vv6)* and *him-3(vv6); cct-4(vv39*) mutants. Germlines were divided into 5 equal sized zones that largely correspond to the following stages in wild-types: premeiotic (Zone 1), transition zone (Zone 2), early pachytene (Zone 3), midpachytene (Zone 4), late pachytene (Zone 5). Pairing levels in all genotypes are not different in the premeiotic region p<0.001, Fisher’s Exact Test), but are reduced in *him-3(vv6)* and *him-3(vv6); cct-4(vv39)* mutants in comparison to wild types in all zones following meiotic entry (p<0.0001). Pairing levels in *him-3(vv6)* and *him-3(vv6); cct-4(vv39)* mutants is not different upon meiotic entry (Zone 2, p=0.81), but rise as pachytene progresses in *him-3(vv6); cct-4(vv39)* in comparison to *him-3(vv6)* (Zone 3, p=0.03; Zone 4, p=0.003, Zone 5 p<0.0001). **B)** Histograms showing percentage of early, mid and late pachytene nuclei with paired and unpaired chromosomes V (5S rDNA locus) in association with the SC marker SYP-1. The region of the germline corresponding to pachytene in wild-types was divided into three equally-sized zones that were largely populated by nuclei of the indicated stages. *him-3(vv6)* and *him-3(vv6); cct-4(vv39)* mutant germlines have extended transitions zones, but eventually show exit from leptotene/zygote stages as evidenced by loss of a polarized nuclear organization and extensive synapsis (not shown). Categories: unpaired and unsynapsed (unpaired FISH signals without SYP-1 tracts); paired and unsynapsed (paired FISH signal without SYP-1 tract), homologously synapsed (paired FISH signals with SYP-1 tract), nonhomologously synapsed and unsynapsed (unpaired FISH signals, one with SYP-1 tract), and nonhomologously synapsed (unpaired FISH signals, both with SYP-1 tracts). In comparison to *him-3(vv6)* mutants, *him-3(vv6); cct-4(vv39)* suppressed germlines have higher levels of homologously synapsed chromosomes (p<0.001 for all stages), fewer participating in nonhomologous synapsis (one or both chromosomes engaged in nonhomologous synapsis; p<0.0001, p<0.0001, p<0.001 in each stage), and an increase in the unpaired and unsynapsed class (p=0.04, p=0.005, p=0.01 at all stages). **C)** Immunofluorescence micrographs showing DNA, HIM-3 and SYP-1 localization in pachytene nuclei in germlines of the indicated genotypes. White arrows indicate regions of unsynapsed chromosome axes marked by HIM-3 without detectable SYP-1. Scale bars, 5 µm.

We next examined the effect of *cct-4(vv39)* on the extensive nonhomologous synapsis that is a characteristic of *him-3(vv6)* mutants, reasoning that the improved pairing observed in the presence of the suppressor should result in an increase in the homologously synapsed chromosome population at the expense of the nonhomologously synapsed one; chromosomes competent for PC protein recruitment and pairing would be stabilized by synapsis while the remainder engaged in nonhomologous synapsis. Nonhomologous synapsis in *him-3(vv6)* mutants was investigated by simultaneously assaying the pairing status of chromosome V using FISH and its participation in synapsis using SYP-1 as a marker of SC formation (Fig. 3B; ^49^). Scoring of the 5S rDNA probe relative to SYP-1 localization was technically unreliable in the tightly clustered chromatin of wild-type TZ nuclei and it was instead assessed in three equally-sized zones corresponding to the early, mid and late pachytene regions of the wild-type germline. In the case of *him-3(vv6)* and *him-3(vv4); cct-4(vv39*) mutants, the assay was technically possible given the partial polarization of nuclei in the extended TZ and truncated pachytene region, and their germlines were divided into 3 regions spatially corresponding to the 3 pachytene regions of wild-type germlines for analysis. In wild types, 87% of early pachytene nuclei had homologously synapsed chromosomes; the remaining population consisted of nuclei with paired and unsynapsed chromosomes that progressed to achieve 98% homologous synapsis by late pachytene. In contrast, only ∼12% of nuclei in any of the regions tested in *him-3(vv6)* mutants exhibited homologous synapsis; however, *cct-4(vv39)* suppressed germlines reached 43% in the earliest stage and rose to 55% by late pachytene. As predicted, the trend towards higher levels of homologous synapsis in *him-3(vv6); cct-4(vv39)* mutants was accompanied by a corresponding decrease in the nonhomologously synapsed populations. In *him-3(vv6)* mutants, the number of nuclei with at least one nonhomologously synapsed chromosome was already 70% in the earliest nuclei assayed and rose to 80% by the end of pachytene. In contrast, only 20% of early prophase nuclei and 25% of late pachytene nuclei in suppressed germlines showed nonhomologous synapsis, indicating that the increased pairing observed in the presence of the suppressor was stabilized by synapsis, leading to a reduction in the nonhomologously synapsed population.

Unexpectedly, our analysis also revealed the presence of significantly higher levels of unsynapsed chromosomes in the suppressed germlines in comparison to *him-3(vv6)* mutants alone (18% and 4%, respectively at late pachytene; p=0.01), a phenomenon accompanied by the appearance of more unpaired tracts of DAPI-stained chromosomes and a visible increase in HIM-3-marked axes lacking SYP-1 (Fig. 3C). These results indicate that *cct-4(vv39)* not only improved pairing in *him-3(vv6)* mutants, but it also reduced SC formation between unpaired chromosomes, an observation that can be explained by 1) the restoration of chromosome movement (which discourages nonhomologous synapsis), and 2) improved HTP-1/2 axis localization (which improves co-ordination of pairing and synapsis). Taken together, our data are consistent with the interpretation that CCT-4^P382F^ improves HIM-3^S35F^ loading, which mitigates the delayed axis morphogenesis defects of *him-3(vv6)* mutants, and in turn results in autosomal PC protein localization, establishment of PC-mediated connections to cytoplasmic forces, and an increase in homologous synapsis and crossing over.

### cct-4(vv39) defines an interface of CCT-4 required for HIM-3 axis localization

We next sought to understand the mechanism of CCT-4^P382S^-mediated suppression of the defect in HIM-3^S35F^. The HORMA domain has a complex tertiary structure that would be predicted to require active assistance in folding; consequently, a plausible scenario is that HIM-3 is a client of the TRiC chaperonin and that the S35F substitution resulting from *vv6* disrupts its interaction with a CCT-4 interface required for its folding into a binding-competent conformation. The P382 residue affected by *vv39* is in the intermediate domain, a region that contributes to substrate recognition and acts as a hinge to open and close the complex during substrate binding (Fig. 1D; ^82,83^), suggesting that P382 is either part of the substrate recognition interface for the mHORMADs and/or positions them for interaction with the interface(s). If CCT-4^P382S^ restores HIM-3^S3F^ function through a physical interaction, the suppression of *him-3(vv6)* loading defects by *cct-4(vv39)* would reflect suppression by a compensatory structural change in CCT-4; the P832S substitution in CCT-4 specifically modifies the interaction interface of the subunit with HIM-3 and physically accommodates the structural change caused by the S35F substitution in HIM-3 during its folding (Model, Fig. 4A). If true, the suppressor activity of *cct-4(vv39)* would be predicted to be 1) dependent on the presence of the *him-3* protein, and 2) restricted to the *vv6* mutation. We tested the basis of the *vv39* suppression and found that *vv39* did not affect the embryonic lethality and X-chromosome nondisjunction of *him-3* null mutants (p>0.001 Kruskal-Wallis and Dunn’s post test), indicating that the suppression required HIM-3. Furthermore, *cct-4 (vv39)*-mediated suppression was specific to the *vv6* allele since *vv39* did not rescue other *him-3* missense alleles with mutations in the HORMA domain *(e1147, e1256, me80;* p>0.001 for all genotypes).

**Figure 4.**
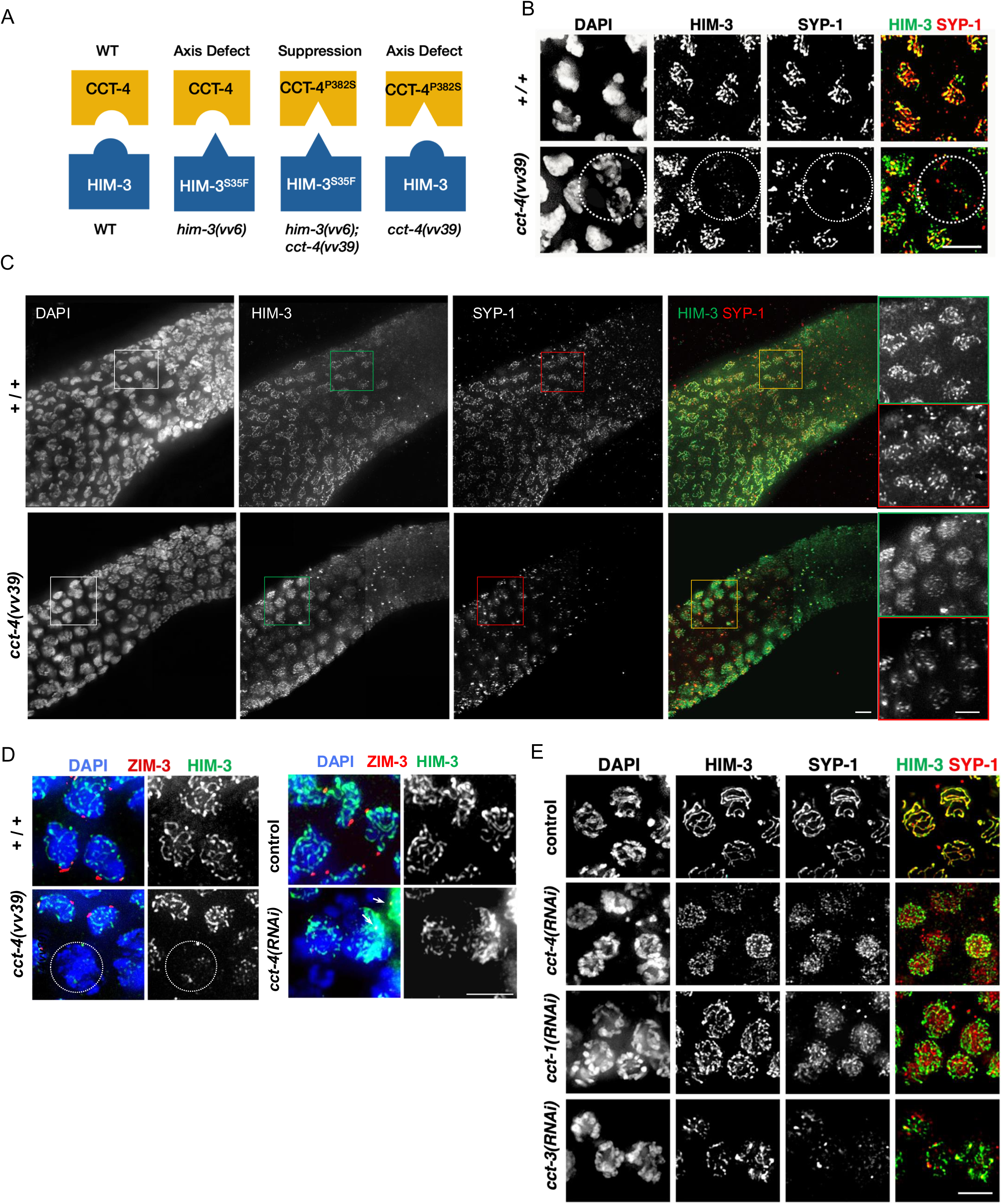
Loss of function of TRiC subunits results in meiotic axis defects. **A)** Model for the interaction of CCT-4 and HIM-3 wild-type and mutant variants. **B)** Immunofluorescence micrographs showing DAPI-stained chromatin and localization of HIM-3 and SYP-1 in early TZ (leptotene/zygotene) nuclei of *cct-4(vv39)* mutant germlines. In *cct-4(vv39)* mutant germlines, the TZ contains intermixed polarized and non-polarized nuclei (circled). HIM-3 and SYP-1 can be detected in nonpolarized nuclei, but axes are immature. **C)** Intact three-dimensionally preserved germlines of wild-types and *cct-4(vv39)* mutants stained as in B) showing HIM-3 and SYP-1 localization. Pachytene zone (bottom left) and proliferating region (upper right). Insets are magnifications of the indicated regions. **D)** Nuclei from the leptotene/zygotene region (HIM-3 positive) of germlines from animals of the indicated genotypes stained for DNA, the PC binding protein ZIM-3, and HIM-3. In *cct-4(vv39)* mutant nuclei, ZIM-3 is localized exclusively to the p e r i p h e r y o f polarized nuclei with robust loading of HIM-3. In *cct-4(RNAi)* germlines, disorganized HIM-3 axes appear, but fail to form bright foci although small, faint foci are occasionally detectable (arrows). **E)** Nuclei from the region corresponding to midpachytene in wild types in empty vector (control) and germlines depleted for the indicated TRiC subunits stained for DNA, HIM-3, and the SC component SYP-1. Because of cell-cycle and oogenesis defects, precise staging of nuceli was not possible (see text). Scale bars, 5 µm.

*cct-4* is an essential gene and the surprising wild-type viability and fertility of *cct-4(vv39)* mutants indicates that the P382S substitution does not detectably affect the folding of other proteins by the chaperonin. However, we observed that *cct-4(vv39*) mutant germlines contained TZ nuclei that exhibited delayed localization of the wild-type HIM-3 protein, loss of ZIM-3 localization, and SYP-1 localization into puncta or short fragments (Fig. 4B-D). These nuclei could occasionally be seen interspersed among nuclei that showed the reverse (robust localization of HIM-3, ZIM-3, and SYP-1; Fig. 4B,D), indicating that the HIM-3 loading defects are cell autonomous. We conclude that *cct-4(vv39)* mutants show an early delay in HIM-3 localization, PC assembly, and synapsis that is ultimately corrected without affecting CO formation and fertility. Importantly, these data indicate that the presence of CCT-4^P382S^ in combination with wild-type HIM-3 resulted in phenotypes associated with axis assembly defects, as predicted by our interaction model (Fig. 4A); although these defects were less penetrant, they nevertheless qualitatively replicated those observed in the reverse combination (CCT-4*^+/+^* and HIM-3^S35F^). Altogether, these results reveal a role for CCT-4 in axis morphogenesis *per se* and suggest that this function requires a physical interaction between the chaperonin subunit and HIM-3.

We next reasoned that if the interaction between mutant CCT-4^P382S^ and wild-type HIM-3 slowed the events surrounding chromosome pairing, *cct-4(vv39)* would be predicted to suppress mutants in which these events were inappropriately accelerated. In *htp-1(gk174)* null mutants that show a premature exit from leptotene-zygotene stages, PC proteins localize and SUN-1 aggregates form in the vicinity of chromosome ends^14,15^, but in the absence of HTP-1, SC assembly is licensed before pairing is accomplished.^21,22^ We observed that *htp-1(gk174); cct-4(vv39)* mutants exhibited significantly less nuclei with 12 DAPI bodies than *htp-1(gk174)* mutants alone (p<0.0001; Fig. 5A), indicating a partial restoration of chiasma formation, and we next assessed the level of pairing. In agreement with previously published results^21,22^, only ∼10% of nuclei at any stage showed pairing of chromosome V in control *htp-1(gk174)* mutant germlines (Fig. 5B). In *htp-1(gk174); cct-4(vv39)* mutant germlines however, pairing levels at leptotene/zygotene increased to 21% in comparison to *htp-3(gk174)* mutants alone (p*=*0.03). By late pachytene, pairing levels in suppressed germlines had reached 29% (p=0.0004) and this was accompanied by a visible increase in HIM-3-marked axes lacking SYP-1 localization (Fig. 5C), consistent with increased pairing at the expense of nonhomologous synapsis. These results indicate that the pairing and nonhomologous synapsis defects of *htp-1(gk174)* mutants can be partially suppressed by *cct-4(vv39*). This phenomenon is most simply explained by the discrete delay in HIM-3 loading in the presence CCT-4^P382S^, which has the practical effect of simultaneously extending the homology search window and suppressing SC formation, giving chromosomes more time to pair and appropriately nucleate the SC at the pairing centers. In aggregate, these results suggest that an interaction between CCT-4 and HIM-3 is required for timely axis assembly and cell-cycle progression.

**Figure 5.**
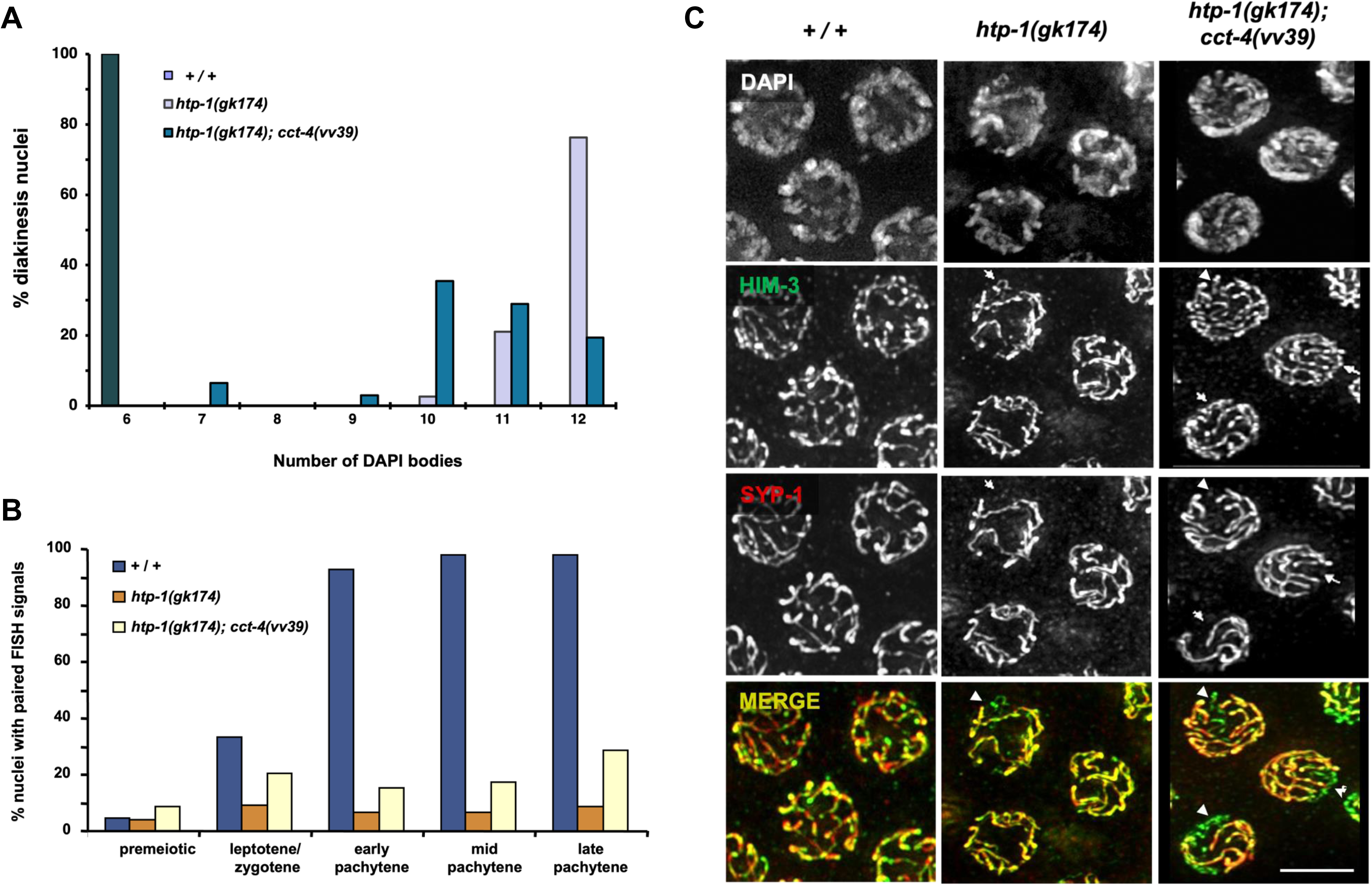
*cct-4(vv39)* partially suppresses pairing and CO formation in *htp-1(gk147)* mutants. **A)** Histogram showing the percentage of diakinesis nuclei at the minus 1 position of the germline (abutting the spermatheca) with the indicated number of DAPI-stained figures. *htp-1(gk174); cct-4(vv39)* mutants have significantly less nuclei with 12 DAPI bodies than *htp-1(gk174)* single mutants (p<0.0001) and more nuclei with 7 and 10 DAPI bodies (p=0.007 and p<0.0001 respectively). **B)** Histogram showing pairing levels in germline nuclei of the indicated stages as assayed using a FISH probe targeting the 5S rDNA locus. Pairing levels are significantly increased in *htp-1(gk174); cct-4(vv39)* mutants at leptotene/zygotene in comparison to *htp-1(gk174)* (p=0.03), but are still very low in comparison to WT (p*=*0.04) at the same stage. The level of pairing attained by late pachytene is also significantly different higher in *htp-1(gk174); cct-4(vv39)* mutants in comparison to *htp-1(gk174)* mutants (p=0.0004). **C)** Immunofluorescence micrographs showing DAPI-stained DNA, HIM-3, and SYP-1 localization in mid-pachytene nuclei. Unsynapsed axes are indicated by white arrows, and correspond to chromosome axes marked by HIM-3 without colocalization of SYP-1. Scale bars, 5µm.

### Mutations in *htp-3* and *htp-1* at sites that correspond to *him-3(vv6)* result in axis morphogenesis defects that are suppressed by *cct-4(vv39)*

The missense mutation encoded by *him-3(vv6)* affects a serine (S35F) near the beginning of the first α helix (αA) of the structural core and is widely conserved among all HORMAD proteins, including HTP-1/2/3 (Fig. 1A,B).^84^ In the closed conformation of an mHORMAD protein, helix αA interacts with the safety belt β-strands; partial unfolding of this helix by Pch2/TRiP leads to disengagement of the safety belt from the HORMA domain core and formation of the active unbuckled conformation^31^. The substitution of S35 with a hydrophobic residue at this position would be predicted to disrupt the ability of HIM-3 to adopt the unbuckled active conformation^42^, consistent with the delayed appearance of the mutant protein at chromosome axes. In *C. elegans,* PCH-2 is dispensable for mHORMAD localization, indicating that formation of the unbuckled conformation is catalyzed by an alternate pathway, and we considered the possibility that TRiC performed this function. In the case of *cct-4(vv39)*, the affected P382 lies in the region of the subunit that hinges when the complex changes from an open to a closed conformation (Fig.1B); a serine substitution at this position would be predicted make the hinge even more flexible in the mutant and/or affect its direct interaction with specific substrates during their folding^83,85^. These observations raise the possibility that the altered CCT-4^P382S^ is more effective than its wild-type counterpart at folding mutant HIM-3^S35F^ into an unbuckled conformation which in turn increases the pool of active HIM-3 that can adopt the closed state and bind to HTP-3. We reasoned that if HIM-3 were a client of TRiC that required a specific CCT-4 interface for its folding, the other mHORMADs would be similarly dependent on this interface to fold into proteins competent for axis localization. To investigate this possibility, the *vv6-*encoded substitution (S35F) was introduced by CRISPR genome editing at the corresponding serine in HTP-3 and HTP-1 (S37F and S38F respectively; Fig. 1A), and the resulting mutants were assayed for defects in axis assembly and the ability of CCT-4^P382S^ to suppress them.

In the case of *htp-3(vv153)* mutants, HTP-3^S37F^ co-localized with meiotic chromosomes in early TZ nuclei, but appeared largely as abundant bright puncta that intermittently formed segments instead of the contiguous linear elements that were present in wild-type nuclei at the same stage (Fig. 6A), indicating that HTP-3^S37F^ binding to the chromosome is compromised. HIM-3 in turn adopted a faint and punctate appearance except for stretches that coincided with robust HTP-3^S37F^ localization, while SYP-1 appeared in puncta, stretches, and nuclear aggregates at a stage when wild-type nuclei showed near complete synapsis (Fig 6A, B). However, these early axis assembly and synapsis defects were effectively corrected as prophase progressed and chromosomes appeared fully synapsed at pachytene. Importantly, 94% of the diakinesis nuclei in *htp-3(vv153)* mutants showed 6 bivalents (Fig. 6C,), indicating that the function of the mutant protein with respect to meiotic DSB formation and HIM-3 recruitment (through which synapsis and CO formation are mediated) was largely intact. In *htp-3(vv153); cct-4(vv39)* suppressed germlines, HTP-3 localization appeared more continuous than in *htp-3(vv153)* mutants and supported timely HIM-3 loading and SYP-1 recruitment (Fig. 6A,B). Furthermore, the modest, but significant increase in embryonic lethality observed in *htp-3(vv153)* mutants was also mitigated in the presence of *cct-4(vv39)* (Fig.6D). These results indicate that the introduction of the *vv6* mutation into *htp-3* results in a delay in the appearance of HTP-3 stretches that can support HIM-3 binding and SC formation. The observation that these defects can be suppressed by *cct-4(vv39)* is consistent with the interpretation that an interaction between the HTP-3 HORMA domain and CCT-4 is required for HTP-3 localization to nascent axes and for timely axis morphogenesis.

**Figure 6.**
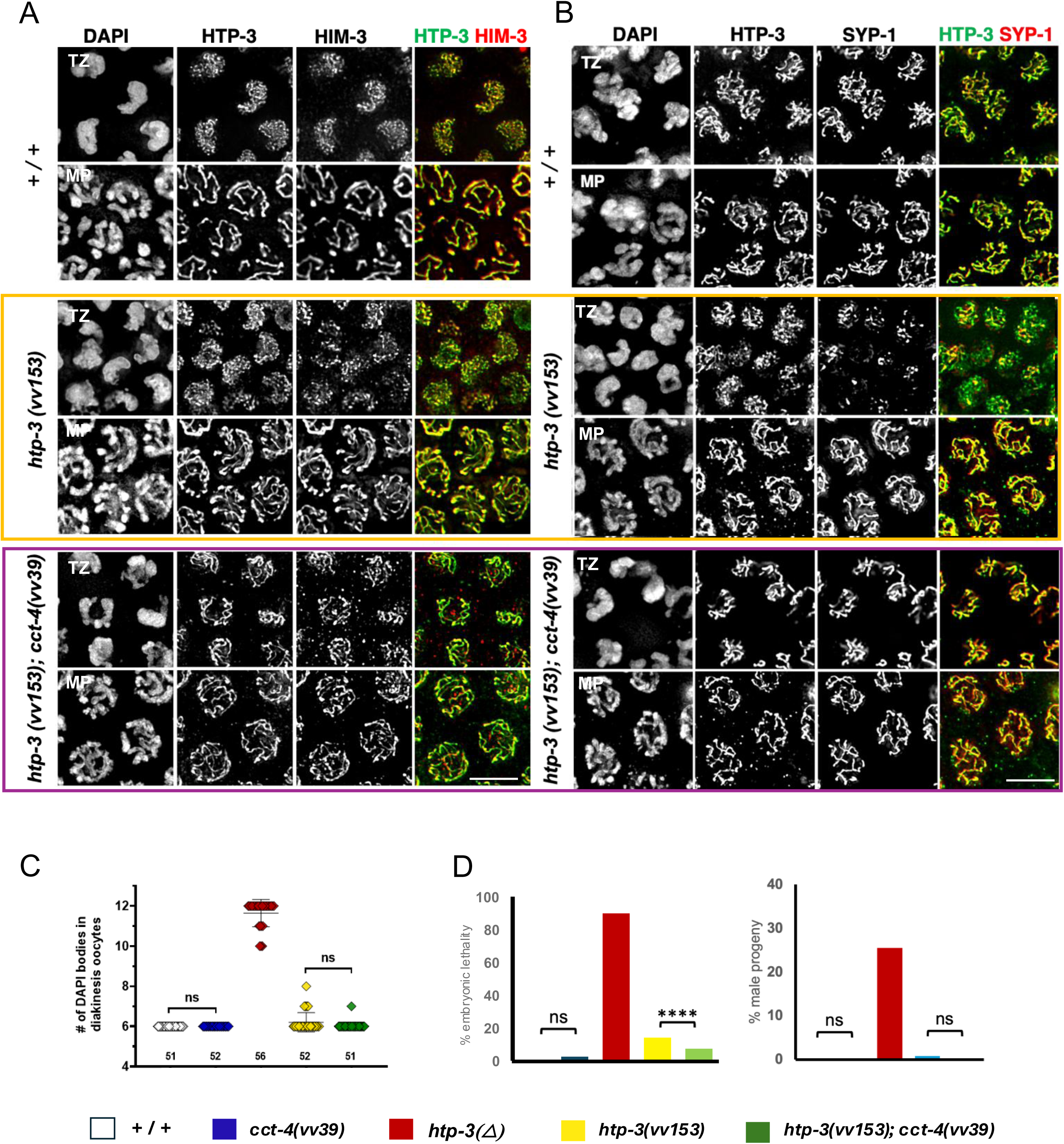
*htp-3(vv153)* HORMAD mutants show delayed HTP-3 loading that is suppressed by *cct-4(vv39)*. Early transition zone and pachytene nuclei from germlines of the indicated genotypes stained for HTP-3 and **A)** and HIM-3 or **B)** SYP-1. **C)** Scatterplot graph showing the number of DAPI-stained bodies in −1 and −2 diakinesis oocytes from germlines of the indicated genotypes. Numbers below data points indicate the number of nuclei scored. The number of DAPI bodies is not significantly different between wild types and the other genotypes using Kruskal-Wallis followed by Dunn’s pairwise comparison test (p>0.05). **D)** Histograms showing the percentage of embryonic lethality and percentage of males among the surviving progeny from hermaphrodites of the indicated genotypes. Chi square test was done on the raw data to measure statistical significance (P value < 0.05 was considered as significant). ns = not significant, **** = P value < 0.000.1. Colors are consistent for each mutant (white is for WT control, blue is for *cct-4(vv39*), red is *htp-3(Δ),* yellow is for *htp-3(vv153)* and green is for *htp-3(vv153); cct-4(vv39)*).

We next turned our attention to HTP-1, which presented a more complex situation given its regulatory functions in chromosome pairing and synapsis. To analyze HTP-1, we introduced the HTP-1^S38F^ substitution in the *htp-2(tm2543)* null mutant background to be able to selectively localize it using immunofluorescence (the antibody detects both proteins; ^45^). In early TZ nuclei of *htp-1(vv164) htp-2(tm2543)* mutants, HTP-1^S38F^ localization to HTP-3-marked axes was incomplete and consisted of numerous small foci of varying intensities that often coalesced, indicating a delay in HTP-1 recruitment (Fig. 7A). This delay was also visibly manifested by delayed synapsis; SYP-1 appeared as foci and 1-2 nuclear aggregates instead of the linear tracts that co-localized with HTP-3 in wild-type nuclei at this stage (Fig. 7B). By midpachytene, however, HTP-1 and SYP-1 localized contiguously along the chromosomes, indicating that HTP-1 levels at the axes increased over time and could eventually support synapsis. However, the eventual recruitment of HTP-1 was not sufficient to support CO formation since the level of univalents in the diakinesis nuclei and the embryonic lethality of *htp-1(vv164) htp-2(tm2543)* mutants did not differ from *htp-1(gk147) htp-2(tm2543)* null mutants (Fig. 7C). To assess the effect of *cct-4(vv39)* on the loading of HTP-1^S38F^ onto chromosome axes, we focused on the early leptotene-zygotene nuclei in the TZ of suppressed mutant germlines; both HTP-1 and SYP-1 appeared earlier and colocalized with the HTP-3-marked chromosomes, indicating that *cct-4(vv39)* partially restored timely loading of HTP-1^S38F^ to the chromosome axis in levels sufficient enough to support extensive SC formation (Fig. 7A,B). Nevertheless, *cct-4(vv39)* did not suppress the CO and embryonic viability defects of *htp-1(vv164) htp-2(tm2543)* mutants (Fig. 7C,D), indicating that the improved HTP-1 localization and SC formation observed in the presence of the suppressor was not sufficient for the timely execution of other meiotic processes required for CO formation, and/or the mutant protein was not capable of functioning in those processes. Collectively, these data are consistent with the model that mutant CCT-4^P382S^ is more effective than its wild-type counterpart at folding mutant HIM-3^S35F^ into an unbuckled conformation that can adopt the closed state and bind to HTP-3. The observation that the binding defects of HTP-3^S37F^ and HTP-1^S38F^ can at least partially be overcome in the presence of CCT-4^P382S^ suggests that the HORMA domain of each paralog is similarly compromised by the *vv6* mutation and can consequently be accommodated by the P392S change in CCT-4 to improve binding. These results reinforce the evidence described above that CCT-4 is required for loading of the meiotic HORMAD proteins during axis assembly at early prophase stages,

**Figure 7.**
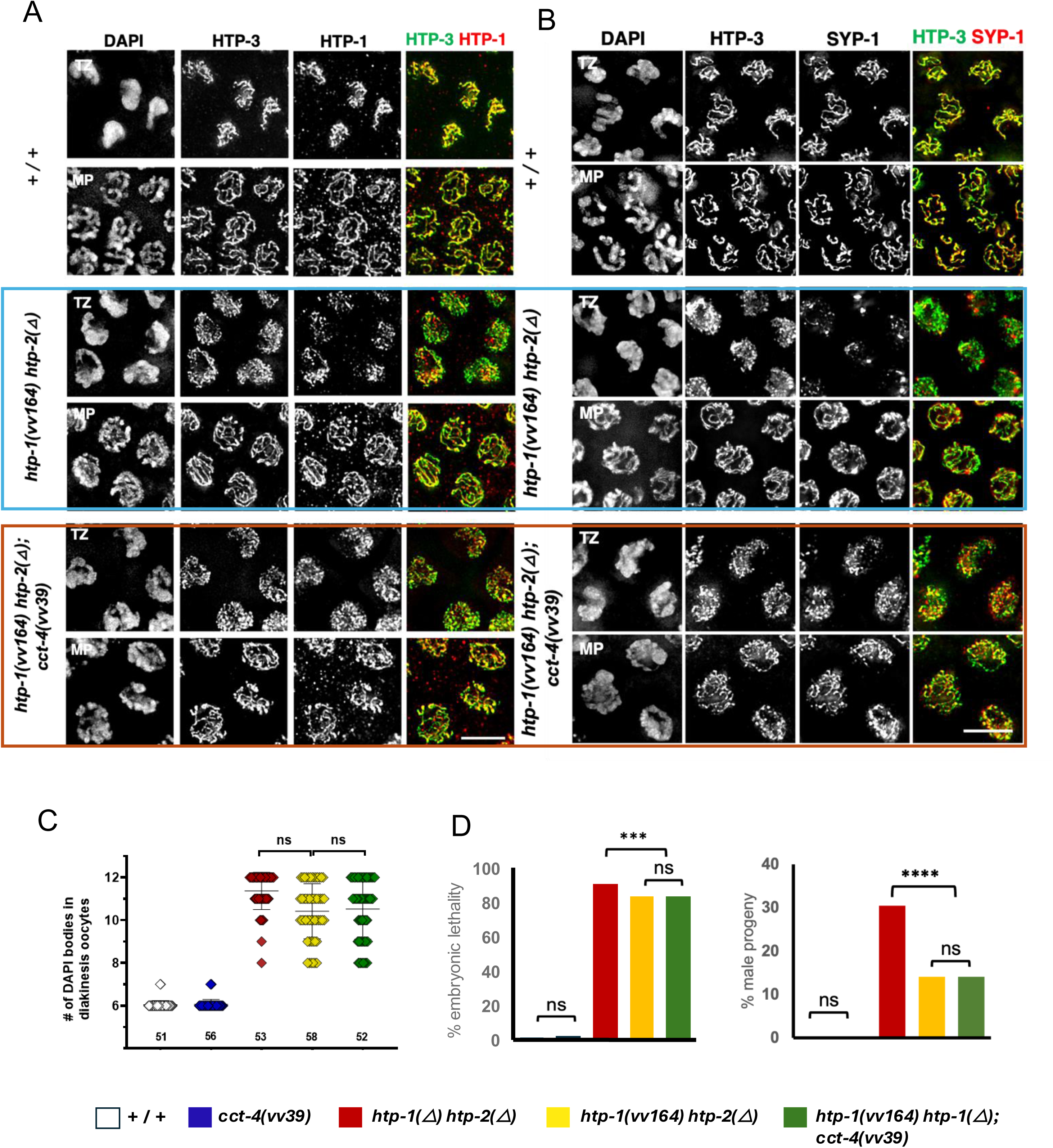
*htp-1(vv164) htp-2(Δ)* HORMAD mutants show delayed HTP-1 loading that is suppressed by *cct-4(vv39)*. Early transition zone and pachytene nuclei from germlines of the indicated genotypes stained for HTP-3 and **A)** HTP-1, or **B)** SYP-1. **C)** Scatterplot graph showing the number of DAPI-stained bodies in −1 and −2 diakinesis oocytes from germlines of the indicated genotypes. Numbers below data points indicate the number of nuclei scored. Using Kruskal-Wallis followed by Dunn’s pairwise comparison test, the number of DAPI bodies in diakinesis nuclei of *htp-1(vv164) htp-2(Δ)* mutants is not different than in *htp-1(Δ) htp-2(Δ)* null mutants and is not suppressed by *cct-4(vv39) (p*>0.05). **D)** Histograms showing the percentage of embryonic lethality and percentage of males among the surviving progeny from hermaphrodites of the indicated genotypes. Chi square test was done on the raw data to measure statistical significance (P value < 0.05 was considered as significant). *htp-1(vv164) htp-2(Δ)* mutants exhibit high levels of embryonic lethality and incidence of males that are not different than *htp-1(vv164) htp-2(Δ); cct-4(vv39)* suppressed germlines. ns = not significant, *** = P value < 0.001 and **** = P value < 0.0001. Colors are consistent for each mutant (white is for WT control, blue is for *cct-4(vv39*), red is *htp-1(Δ) htp-2(Δ),* yellow is for *htp-1(vv164) htp-2(Δ)* and green is for *htp-1(vv164) htp-2(Δ); cct-4(vv39)*).

### *cct-4(RNAi)* results in axis morphogenesis, PC protein recruitment, and synapsis defects

In contrast to lethality associated with null mutations in *cct* genes, *cct-4(vv39)* is a hypomorphic allele that resulted in early disruptions to axis formation that were ultimately corrected as prophase progressed. Previous studies have observed that RNAi depletion of individual *cct* genes results in variable and pleiotropic effects including embryonic lethality, sterility, and defects in gonad morphogenesis and oogenesis; these can be attributed to the role of TRiC in folding substrates required for microtubule dynamics ^86–88^, cell-cycle progression^89^, and regulation of ribonucleoprotein complexes during oogenesis ^90,91^. To further investigate the function of *cct-4* in meiotic prophase, we identified *cct* partial RNAi conditions in which cell proliferation in the mitotic region of the germline was disrupted (Fig. S2; Materials and Methods), but HIM-3 as an indicator of meiotic entry and axis formation could still be detected. *cct-4(RNAi)* germlines were smaller in comparison to controls and had fewer nuclei, with irregularly spaced macro- and micronuclei appearing throughout (Fig. S2), consistent with loss of TRiC activity in folding substrates required for germ cell divisions and oogenesis. In the region of the *cct-4(RNAi)* germline spatially corresponding to the TZ of control germlines, nuclei lacked the polarization of chromatin characteristic of this stage was lost, but HIM-3 could be detected on chromosomes, indicating that the nuclei had entered meiotic prophase. In these nuclei HIM-3 localization was highly disrupted, appearing as variably-sized segments and foci (Fig. 4 B,C), reminiscent of the immature chromosome axis morphology observed in *cct-4(vv39)* mutants (Fig. 4A). Furthermore, ZIM-3 was not localized to the nuclear periphery, although a small focus could occasionally be detected, indicating that the protein was expressed (Fig. 4B). In more proximal nuclei located in the pachytene region, the discontinuous appearance of HIM-3-marked chromosome persisted, and SYP-1 remained diffuse in the nucleoplasm (Fig. 4D). These results indicate depletion of *cct-4* disrupts HIM-3 localization and the formation of axes that can support PC binding protein recruitment and SC formation. Taken together, these data confirm that CCT-4 is essential for meiotic chromosome axis morphogenesis and SC assembly.

### Loss of CCT-1 and CCT-3 recapitulates *cct-4(RNAi)* axis morphogenesis defects

While it is well-established that each CCT subunit is essential for the formation and function of TRiC, examples of complex-independent functions of some subunits have been reported^92^. The *C. elegans* genome encodes eight CCT subunits (CCT-1 through 8) that are orthologs of the conserved eight subunits of TRiC^93,94^. To determine if the axis morphogenesis function of CCT-4 occurs in the context of the chaperonin, we reasoned that germline disruption of other subunits of the complex should mimic the effect of *cct-4(RNAi)*. Depletion of *cct-1* or *cct-3* by RNAi resulted in meiotic defects that resembled those observed in *cct-4(RNAi)* germlines; the HIM-3-marked axis morphology of nuclei in the region of the germline corresponding to pachytene appeared immature and disorganized, and SYP-1 was diffuse within the nucleus (Fig. 4D). Since loss of each of three essential subunits of TRiC independently resulted in defects in axis morphogenesis and SYP-1 recruitment, we conclude that the simplest explanation for these results is that meiotic axis assembly requires the chaperonin complex and its protein folding function.

### CCT-4 localizes to germline nuclei and colocalizes with chromatin during meiotic chromosome pairing stages

While TRiC has well established roles in conserved cytosolic processes in *C. elegans*, previous developmental studies have noted nuclear localization of CCT-1 and CCT-5^68,95^. Since the requirement for *cct-4* in meiotic chromosome assembly suggests a nuclear function for the protein in the germline, we examined the localization of CCT-4 using antibodies raised against a 16 a.a. peptide unique to the CCT-4 subunit (Fig. 8A) and confirmed their specificity in *cct-4* depleted germlines (Fig. S3). Consistent with its function as a subunit of TRiC, CCT-4-marked foci were abundant throughout the cytoplasm of wild-type germlines. During meiotic prophase, CCT-4 was also detected as a bright punctate nuclear signal that also showed broad colocalization with chromatin when observed through the nuclear volume; TZ nuclei showed heterogenous levels of CCT-4 foci that were grossly enriched in the vicinity of the chromosomes, and upon pachytene entry nuclear CCT-4 levels dramatically increased and persisted until dispersing into the nucleoplasm at diakinesis (Fig. 8A). These observations indicate that CCT-4 localizes to meiotic prophase nuclei and suggest the presence of a nuclear population of TRiC that associates with chromosomes during pairing and SC assembly stages.

**Figure 8.**
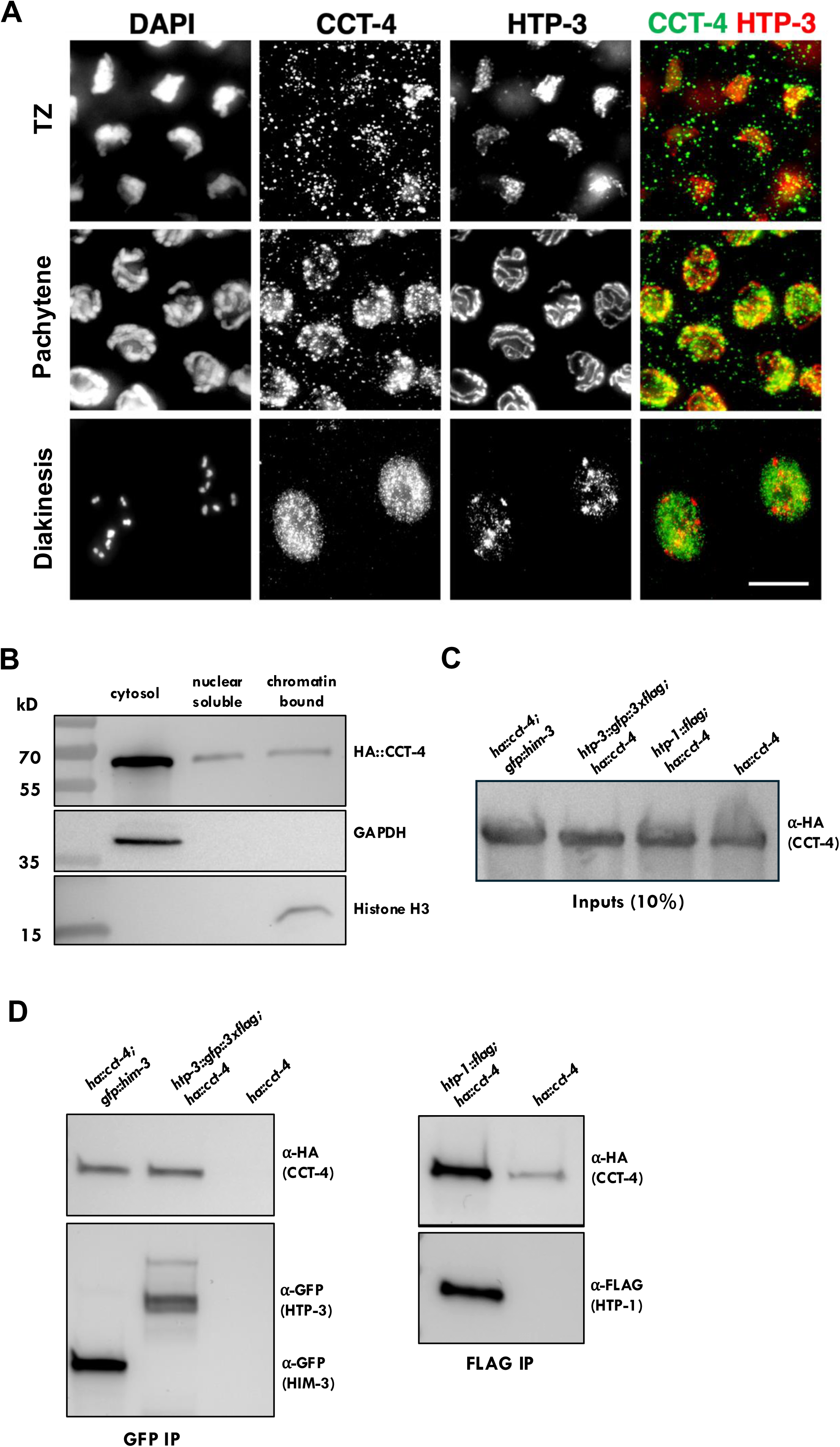
CCT-4 localizes to meiotic prophase nuclei and forms complexes with mHORMADs *in vivo.* **A)** Wild-type germlines stained with DAPI, α-HTP-3 and α-CCT-4 in nuclei of the indicated meiotic stages. **B)** Western Blot analysis of fractionated protein extracts shows that HA::CCT-4 is enriched in the cytoplasm, as well as present in comparable amounts in both nuclear-soluble and chromatin-bound fractions. GAPDH and histone H3 were employed as loading controls for cytosolic and nuclear insoluble fractions respectively. **C)** Western blot analysis showing abundance HA::CCT-4 in fractions used for the inputs for the immunoprecipitation. Equal amounts of protein extracts were loaded. **D)** HA::CCT-4 forms complexes with meiotic HORMAD proteins *in vivo*. Left: Western blot analysis reveals that both GFP::HIM-3 and HTP-3::GFP co-immunoprecipitate with HA::CCT-4. Right: HA::CCT-4 physically interacts with HTP-1::FLAG.

### mHORMADs form nuclear complexes with TRiC subunits

The model that a CCT-4 interface interacts with HIM-3 during axis assembly necessarily demands a physical interaction between the two, and we next sought to confirm the nuclear presence of CCT-4 and whether HIM-3 and CCT-4 form a complex *in vivo*. Using functional endogenously-tagged proteins, we first immunoprecipitated HA::CCT-4 from cytosol, nuclear soluble, and DNA-bound protein fractions ^96^. Given its essential role in the chaperonin complex CCT-4 was predictably abundant in cytosolic extracts, however, it was also easily detectable in nuclear extracts and appeared at similar levels in nuclear soluble and nuclear insoluble fractions containing chromatin (Fig. 8B). These results converged with our localization data that detected nuclear CCT-4 that partially overlapped with meiotic chromatin. To determine if HIM-3 is found in CCT-4 complexes, CCT-4 was immunoprecipitated from nuclear extracts derived from worms co-expressing HA::CCT-4 and GFP::HIM-3; western blot analysis revealed abundant levels of CCT-4 in HIM-3 immunoprecipitates, consistent with an interaction between the two proteins in germline nuclei (Fig. 8D). Furthermore, CCT-4 also appeared in HTP-1 and HTP-3 immunoprecipitated complexes, demonstrating that the interaction was not HIM-3 specific, but extended to the other meiotic HORMAD proteins (Fig. 8D). We next investigated if the CCT-4 interaction was specific to the CCT-4 subunit, or occurred as part of the TRiC complex, using a functional endogenously tagged *cct-5*, and extracts derived from worms co-expressing HA::CCT-4 and FLAG::CCT-5. Like CCT-4, CCT-5 predictably appeared at high levels in the cytosol, but was also present in soluble and chromatin-bound nuclear fractions at comparable levels to CCT-4 (Fig S4A). Both subunits appeared in the nuclear fractions at similar levels, suggestive of the stoichiometry expected from the structure of the TRiC complex and CCT-5 was detected in immunoprecipitated CCT-4-containing complexes, results that in aggregate indicate the presence of a nuclear TRiC that is tethered to chromatin (Fig. S4B). Finally, CCT-5 could be detected in immunoprecipitated GFP::HIM-3 complexes isolated from worms co-expressing FLAG::CCT-5 (Fig. S4B), indicating that a population of HIM-3 is in complexes with another TRiC subunit, an observation most simply explained by HIM-3 being a substrate of an assembled nuclear TRiC. In summary, our collective results are compatible with interpretation that HIM-3, HTP-3, and HTP-1/2 physically interact with a nuclear TRiC chaperonin, an engagement that reflects a requirement for local protein folding in assembling meiotic chromosome axes.

## DISCUSSION

### A nuclear function for TRiC/CCT in meiotic chromosome organization

Genetic suppressor screens are a classic approach to identifying new factors or pathways functioning in a process of interest. In this study, our screen for genetic suppression of *him-3(vv6*)-associated meiotic defects identified an unexpected role for TRiC complex in meiotic chromosome axis morphogenesis. We isolated a unique allele of *cct-4* that was able to suppress the axis localization defects of the mutant protein encoded by *him-3(vv6)* and partially restore homolog pairing, synapsis, and CO formation in *him-3(vv6)* mutants. The suppressor activity of *cct-4(vv39)* was specific to the *vv6* allele, a phenomenon typical of suppression that is based on structural accommodation of a mutant protein by a corresponding structural change in its interacting partner. This scenario also explains the delayed axis morphogenesis observed in *cct-4(vv39)* mutants; in this case, the wild-type HIM-3 and mutant CCT-4^P382S^ combination would be predicted to have similarly structurally compromised interactions that would interfere with HIM-3 localization. *him-3(vv6)* affects a highly conserved serine in the HORMA domain and introducing this change into the HORMA domains of HTP-3 and HTP-1 also resulted in a delay in their axis localization, demonstrating that the substitution disrupts a common structural feature of the mHORMAD proteins that is required for their localization. The observation that CCT-4^P382S^ was also able to suppress the axis localization defects of HTP-3^S37F^ and HTP-1^S38F^ provides strong genetic evidence that the mHORMADs physically interact with CCT-4, an interpretation corroborated by biochemical evidence demonstrating the presence of the mHORMADs in CCT-4 complexes *in vivo*. In addition to its predicated cytosolic localization, CCT-4 grossly colocalized with chromatin during meiotic prophase and CCT-4 and CCT-5 were abundant in the chromatin and soluble nuclear fractions of germline extracts. These results suggest that a nuclear population of TRiC associates with meiotic chromosomes, similar to previous observations in rat spermatocytes ^69^.

Although CCT-4 (and all other subunits) are essential components of TRiC^68,91,97–99^, some complex-independent functions for individual subunits have been demonstrated in other systems, notably CCT2 as an aggrephagy receptor^92^. However, the presence of CCT-5 in immunoprecipitated HIM-3 complexes (in addition to CCT-4) is a strong indicator that the interaction between the mHORMADs and CCT-4 occurs in the context of an assembled TRiC complex *in vivo*. This is further supported by the observation that RNAi of individual subunits of TRiC resulted in HIM-3 localization defects that varied in their severity, but nevertheless uniformly failed to support SC formation, and in the case of *cct-4*, PC binding protein recruitment, two indicators of functional axis assembly. We attribute the presence of HIM-3-marked axes under these partial RNAi conditions to the presence of a pre-existing population of active TRiC in meiotic nuclei that enabled axis formation to varying degrees. This interpretation is supported by the variable penetrance of SC defects observed in *pas-3(RNAi)* germlines depleted for a subunit of another ubiquitous cytoplasmic complex, the 26S proteosome^100^. Consequently, we conclude that it is the protein folding activity of the chaperonin complex itself, rather than potential TRiC-independent activities of individual subunits that is required for axis assembly.

### CCT-4^382S^ identifies an interacting interface with the mHORMADs required for formation of a binding-competent form

The conserved serine affected by *vv6* is positioned near the beginning of the first α helix of the structural core of the *him-3* protein (Fig. 1A), and the substitution with phenylamine at this position would be predicted to affect the structural integrity of the HORMAD core^42^. Computational modelling of the structure of mutant HIM-3^S35F^ was inconclusive and yielded low confidence structures, however, several lines of evidence suggest that the defect in HIM-3^S35F^ lies in forming the active unbuckled conformation that can bind to HTP-3. First, the mutant HIM-3^S35F^ protein is delayed in its localization into contiguous stretches in early prophase, but uninterrupted HIM-3-labelled axes eventually appear, indicating that the protein has a reduced ability to localize to the axis, but once localized is stably associated. The possibility that the S35F substitution affects the formation of an active conformation of HIM-3 is further supported by the fact that the same disruption to the HORMA domains of HTP-3^S37F^ and HTP-1^S38F^ results in similarly disrupted localization dynamics, a phenomenon that is most simply explained by a common defect in forming the active conformation required for binding to interaction partners at the axis. Importantly, HIM-3^S35F^ can perform its wild-type functions in synapsis and in establishing the barrier to using the sister as a repair template during HR (^49^; this study), indicating that the protein is functional once localized and that the key defect is in the early association of the protein with the axis. In considering the collective evidence, we conclude that TRiC containing the mutant CCT-4^P382S^ subunit is more effective than its wild-type counterpart at folding mutant HIM-3^S35F^ into an unbuckled conformation that can adopt the closed state and bind to HTP-3.

The P382 residue affected by *vv39* resides in the intermediate domain, a region that hinges when the complex changes from an open to a closed conformation ^82^ and that can also interact with substrates during protein folding^83,85^. This suggests that P382 identifies a region in CCT-4 that contacts the mHORMADs during protein folding or positions them for interactions with substrate recognition interfaces in other regions of the central cavity of TRiC. The idea that specific contacts between the mHORMADs and CCT-4 are required for their folding has gained traction from recent studies that resolved the progressively folded states of β-tubulin within the closed TRiC chamber, leading to the model that TRiC provides a topological environment that steers folding of a protein through stepwise formation of partially folded intermediate^83,101^. Based on the interactions of tubulin with the inner wall of the TRiC chamber, this mechanism requires specific contacts with TRiC subunits as each intermediate transforms into the next – by extension implying co-evolution between TRiC and some of its substrates^83,101^. In this context, the observation that *cct-4(vv39)* can specifically disrupt HIM-3 axis localization without affecting the fertility or viability of the mutants can be reconciled; CCT-4 contains an interface tailored for contact with mHORMADs during their folding and this is disrupted in *vv39*, leading to altered folding of the axis components while leaving folding of other essential substrates intact. Conversely, the altered interface in CCT-4^P382S^ partially restores its interaction with the mutant mHORMAD variants and supports their folding into an active conformation for binding during axis assembly (Fig. 4A). To our knowledge these data provide the first functional evidence for specific contacts within the TRiC chamber during protein folding.

### Chromosome-tethered TRiC locally remodels mHORMADs for axis assembly

The TRiC/CCT complex exists in the cytoplasm, raising the question of why the meiotic HORMADs are not folded there into the active unbuckled conformation and imported into the nucleus. HIM-3 and HTP-1/2 bind to HTP-3 *in vitro*^29^, indicating that that they can form stable complexes without the participation of nuclear factors. Consequently, the cytosolic folding of the HORMADs into their active (binding competent) conformation could result in aggregation of the proteins and loss of their function. This problem extends into the nucleus as evidenced by mHORMAD dynamics in cohesion-defective mutants; in these cases, HTP-3 fails to localize to the chromosomes and is instead found in one or two nuclear aggregates that contain HIM-3, HTP-1/2, SC components, and cohesin subunits^102,103^. Moreover, the localization of HIM-3 into the aggregates is dependent on HTP-3, indicating that mHORMAD proteins can ectopically interact *in vivo* and form stable complexes without the involvement of chromosomes^46,47^. *C. elegans* is unique among other meiotic systems in that it has multiple mHORMAD proteins that directly interact. Given that HIM-3, HTP-1/2 and HTP-3 are present in meiotic prophase nuclei from the earliest stages, an outstanding question is how unlicensed binding between them is prevented while interactions at the chromosomes are simultaneously licensed.

Meiotic HORMADs are thought to initially adopt an inactive “self-closed” form in which their HORMA domains engage their own C-terminus; this interaction is released as part of the conformational changes leading to binding of interacting partners^32,50,104^. Although the structure of HTP-3 has not been resolved, we favour the model that nucleoplasmic HTP-3 exists in the self-closed form by interacting with its C terminus, a conformation that would sequester the CMs from interactions with the other mHORMADs. Our results indicate that the *htp-3(vv153)* mutation disrupts HTP-3 localization, providing the first evidence that the HTP-3 HORMA domain is required for its association with chromosomes. Consequently, binding of the HTP-3 HORMA domain to its interacting partner at the nascent axis would necessitate release of the C-terminal tail, and the CMs would become spatially available for binding by HIM-3 and HTP-1/2. This simple binary mechanism could ensure that the mHORMAD protein interactions only occur in the context of axis assembly and identifies the formation of the unbuckled HTP-3 species as the key event.

In budding yeast and plants, the remodelling activity of Pch2/TRIP13 is needed to convert the self-closed mHORMADs into an unbuckled conformation to promote their association with chromosomes^104,105^. In *C. elegans* however, PCH-2 is dispensable for mHORMAD localization^57^, and our results suggest that the TRiC complex performs the remodelling of the mHORMADs into the active form that can interact at axes. Key among these is the observation that mHORMADs containing the serine substitution defined by the *vv6* mutation showed defects in localizing to chromosomes, consistent with the structural prediction that this substitution in the N-terminal helix of the HORMA domain would interfere with the transition to the unbuckled conformation. The fact that *cct-4(vv39)* could partially rescue these loading defects provides strong evidence that TRiC is required for formation of the active unbuckled conformation. Our results are compatible with the scenario that nucleoplasmic pools of the meiotic HORMADs exist in a self-closed conformation that is inactive but ready for assembly into the axis once “unbuckled” by chromosome-associated TRiC. A situation with parallels has been described for TRiC-mediated control of the assembly of functional HDAC complexes; inactive HDAC1 is delivered to a nuclear TRiC complex to complete folding of a β sheet before its release in its active form and incorporation into HDAC complexes with correct timing^106^. This has led to the proposal that TRiC can cage proteins in an assembly competent, but inactive state prior to their inclusion into complexes, in this case to prevent uncontrolled deacetylase activity by nascent HDAC1 outside of the regulatory confines of the HDAC complex. We speculate that TRiC-mediated folding similarly addresses the issue of uncontrolled binding activity ouside of the confines of axis assembly, in this case by creating a spatially-confined pool of active mHORMADs where axis assembly is occurring. An appealing aspect of such compartmentalized just-in-time protein folding is the ability to choreograph complex processes by regulating the folding of essential actors into their active form at the time and place that they are needed^107^. Applied to the mHORMADs, chromosome-tethered TRiC could similarly hold self-closed inactive forms of the mHORMADs prior to releasing them in their unbuckled form to interact exclusively in the context of axis assembly. A spatially restricted activity during meiosis has also been observed for the 26S proteosome, which shows conserved localization to meiotic chromosomes, and is essential for events leading to pairing, synapsis, and CO formation^100,108^; in this case an example of local protein degradation as a mechanism of regulating meiotic prophase processes. In summary, our study provides evidence that folding of the *C. elegans* mHORMAD proteins into an active conformation by the TRiC chaperonin is essential for axis assembly and downstream meiotic processes. Moreover, we contribute to the growing evidence of nuclear functions for the chaperonin and suggest that localized protein folding may be an unappreciated and more widely conserved mechanism of regulating meiotic chromosome dynamics and other nuclear processes.

## Acknowledgments

This study is based on the PhD thesis of Dr. Kalun Law, who was supervised by M.Z. during his doctoral work. After exhaustive searches, we were not able to locate him and receive the editorial approval of the manuscript necessary to include him in the authorship and assume his rightful place as first author. The authors would like to acknowledge the late Dr. Jason Young (McGill University) for his infectious enthusiasm for protein folding and his support of this project. We would also like to thank past and current members of the Zetka lab for feedback and helpful discussions, and Ryan Dawson for technical assistance. Some strains were provided by the CGC, which is funded by NIH Office of Research Infrastructure Programs (P40 OD010440), and the *C. elegans* Gene Knockout Consortium. We would also like to thank Yuji Kohara and the Sanger Center for strains and clones; and Monica Colaiacovo, Verena Jantsch, Enriquez Martinez-Perez and Anne Villeneuve for antibodies. This work was supported by the Czech Science Foundation (grant GA23-04918S to N.S.), the Canada Research Chairs program (S.J.J), and the Canadian Institutes of Health Research (grant PJT-173381 to M.Z.).

## Experimental Procedures

### *C. elegans* genetics and culture conditions

*C. elegans* strains were maintained on *E. coli* (OP50)-seeded NGM agar plates at 20°C under standard conditions^109^. All strains were generated in an N2 Bristol background unless otherwise indicated. A full description of the strains used in this study is provided in Supplemental Table 1.

### EMS mutagenesis/screening

EMS (ethyl methane sulfonate) mutagenesis was performed according to standard procedures ^109^. Briefly, *him-3(vv6) unc-24(e138)* hermaphrodites at the L4 larval stage were treated with 25 mM EMS (Sigma) in M9 for 4 h at 20 °C. Worms were washed 4X in M9 media and plated onto seeded NGM plates. F1 were individually plated and their F2 progeny individually cultured and screened for the presence of >20 progeny in the F3 generation. Candidates were outcrossed three times with N2 males to eliminate background mutations.

### Scoring of DAPI bodies and transition zone length

The DAPI-body content of the −1 oocyte was analyzed individually using 3D reconstructed images in Zen application (Blue edition, Version 3.7). A DAPI-body was defined as any continuous DAPI-stained chromatin structure, irrespective of its size. Transition zone length was scored in intact DAPI-stained germlines by measuring the distance from the first distally positioned to the last proximally positioned nucleus showing a polarized conformation.

### Cytological Preparation of gonads and staining

Hermaphrodite gonads were dissected 22-24h post-L4 stage in 1xPBS and fixed with 1% paraformaldehyde for 5 minutes. Samples were frozen in liquid nitrogen then placed in methanol at −20°C for 1 minute. Slides were washed three times with 1xPBST (1xPBS, 0.1% Tween-20) and were blocked in 1% BSA in 1xPBS for 1 hour. The primary antibodies were applied overnight at 4 °C. Antibodies were diluted in 1xPBS with 1% BSA as follows: rabbit **α**-CCT-4 1:50 (this study), rabbit **α**-HIM-3 1:200 ^48^, guinea pig **α**-HTP-3 1:700 ^23^, rabbit **α**-HTP-3 1:200, rabbit **α**-HTP-1/2 1:300, rabbit **α**-HIM-8 1:500 (Novus Biologicals, #41980002), rabbit **α**-PLK-2 1:10 ^80^, rabbit **α**-RAD-51 1:200 ^110^, rabbit **α**-RAD51 1:2000 (Novus Biologicals, #29480002), guinea pig **α**-SYP-1 1:800 ^77^, guinea pig **α**-ZIM-3 1:100, rabbit **α**-GFP 1:2000, guinea pig **α**-pSUN-1 (against serine 12) 1:1300 (gifted by Dr. Verena Jantsch), rabbit **α**-SYP-1 1:1000 (gifted by Dr. Nicola Silva), rabbit **α**-MAD-2 1:10000 ^111^ (gifted by Dr. Arshad Desai), mouse **α**-GFP 1:200 (Abcam, ab291), mouse **α**-FLAG 1:500 (Sigma Aldrich F1804) and mouse **α**HA 1:100 (Cell Signaling Technology, #2367). After three washes in 1 X PBST, secondary antibodies were applied for 2 hours at room temperature. Secondary antibodies that was used in this study are as follows: Alexa Fluor 488 goat anti-rabbit IgG 1:1000 (Invitrogen, A11034), Alexa fluor 488 Donkey anti-guinea pig IgG 1:600 (Jackson ImmunoResearch #166451), Alexa Fluor 488 goat anti-guinea pig IgG 1:1000 (Invitrogen A-11073), Cy3 donkey anti-rat IgG 1:300 (Jackson ImmunoResearch #55721), Alexa Fluor 555 goat anti-rabbit IgG 1:1000 (Invitrogen, #A21429), Alexa Fluor 555 goat anti-guinea pig IgG 1:1000 (Thermo Fisher A-21435), Alexa Fluor 488 donkey anti-mouse IgG 1:500 (Jackson ImmunoResearch #106498). After three additional washes with 1 X PBST, samples were mounted with Vectashield antifading medium (Vector Laboratories, Burlingame, CA) containing 2 µl/ml 4’,6-diamidino-2-phenylindole (DAPI).

### Fluorescence In Situ Hybridization (FISH)

For FISH analysis, dissection, fixation, and hybridization of the gonads were essentially performed as described in ^112^, with the following modifications: worms were dissected in PBS (137 mM NaCl, 2.7 mM KCl, 10 mM NaHPO4, and 2 mM KH2PO4), and the gonads were fixed in 3.7% paraformaldehyde (Electron Microscopy Sciences) in PBS for 1 min, treated with RNase (200 μg/ml, 1 hr at 37°C), and denaturated at 92°C for 90 s. A PCR-amplified 5S rDNA repeat was used as probe for the right end of chromosome V ^76^. The probe was labeled with digoxigenin-11-dUTP (Roche) and the hybridized probe was detected with Cy3-conjugated anti-digoxigenin antibody (1:100; Jackson ImmunoResearch).

### FISH and anti-SYP-1 staining for evaluation of nonhomologous synapsis

FISH was first performed with the 5S rDNA probe as above. Instead of adding the Cy3-conjugated anti-digoxigenin antibody, anti-SYP-1 was applied overnight at 4 °C. After four washes with 1 X TBST-BSA (1 X TBS, 0.1% Tween 20, 1% BSA), Cy3-conjugated anti-digoxigenin antibody and anti-guinea pig-Alexa 488 were applied for 2 hours at room temperature. After three more washes with 1 X PBST, the samples were mounted with DAPI in Vectashield as above.

### Time-course analysis of pairing

Data for three complete gonads were collected for each genotype and/or each probe used. For each gonad, stacks of 30 optical sections were collected in increments of 0.2 µm covering the entire volume of one layer of nuclei, and an image of the entire gonad was assembled using Photoshop 6.0. The region extending between the first mitotic nuclei and the last pachytene nuclei was divided into five equally sized zones. FISH signals were then scored by examination of each single nucleus through its volume; signals were considered paired if the distance between the signals was <0.7 µm ^81^. Data for each zone of the three gonads were pooled together, giving a total number of nuclei with paired/unpaired signals. In the case of simultaneously assaying pairing and synapsis, the status of chromosome V alignment was first ascertained (paired or unpaired FISH signals), and then the DAPI-stained chromosome region to which the FISH signal mapped was examined for SYP-1 colocalization.

### Image acquisition and processing

Images of z-stacks at 0.2μm intervals were acquired using a 100X objective lens using a Delta Vision Deconvolution system equipped with an Olympus 1X70 microscope or a 63X objective lens of a wide-field microscope (Zeiss Axio Observer inverted microscope platform). Images acquired by Delta Vision microscope were processed using Fiji ImageJ^113^ to make a 2D maximized Z projected image for all the specimens except for MDF-2 slides in which a partial Z-projection was done^111^. Z projected images processed by Adobe Photoshop (version 23.0.2). Images acquired by wide-field microscope processed by Zeiss Zen (blue edition, version 3.7.97.04000) to make maximized or partial Z projected 2D images and then deconvoluted. The final images processed and made by Adobe Photoshop (version 25.12.0). For each staining at least 20 germlines were analyzed.

### Production of Antibodies

Mouse polycolonal antibody against CCT-4 was generated using synthetic peptide CKIINSENDSNVNLKM according to standard protocol (Genescript Corporation, Piscataway, NJ) and purified by immunoaffinity chromatography with activated immunoaffinity supports (Affi-Gel 10, Bio-Rad) according to the manufacturer’s instructions. The specificity of the antibody was verified by failure to detect immunofluorescence staining signals in *cct-4(RNAi)* germlines (Fig. S3).

### Quantification of DAPI bodies, embryonic lethality and higher incidence of males

The number of DAPI-stained bodies at diakinesis was scored in the minus 1 and 2 oocytes (most proximal) of dissected germlines and their distribution was statistically assessed using a Kruskal-Wallis test followed by Dunn’s multiple pairwise comparison tests. Embryonic lethality and incidence of males were statistically tested by one-way ANOVA followed by multiple pairwise comparison tests. All calculations were performed by Prism 8 (GraphPad) and p values less than 0.05 were considered significant.

### RNA interference

RNA interference experiments were performed as described in ^114^ using the following oligos for dsRNA templates amplification:

*cct-1:* 5’-TAATA CGACTCACTATAGGAAACGCCGCATTGATAAAAT-3’ and 5’-TAATACGACTC ACTATAGGGGGATCAATTGAGCTTTGGA-3’

*cct-3:* 5’-TAATACGAC TCACTATAGGAATCATGCGTGAAGAGGACA-3’ and 5’-TAATACGACTCACT ATAGGCCTTTGATGGTCCACGAAGTA-3’

*cct-4:* 5’-TAATACGACTCACTATAGGAGCCAGAGAGTGTTCGCAAT-3’ and 5’-TAATA CGACTCACTATAGGCTTCAACCGTGTCTCCCATT-3’

To investigate the loss-of function phenotypes of *cct-1, 3*, and *4*, we depleted each gene by introducing RNAi by as previously described for *htp-3(RNAi)* ^23^. Staged adult hermaphrodites (20–22 hr post-L4) were injected with ∼500 ng/μl dsRNA in the pachytene region of each gonad arm and transferred to fresh plates every ∼15 hr. At 70 hr post-injection, the worms were dissected, fixed, and stained as described above.

### *In-vitro* Mutagenesis

The *cct-4(vv39)* mutation was generated by *in-vitro* mutagenesis using the PCR-based Site-Directed Mutagenesis System kit (Invitrogen), and the mutagenesis primers 5’-AGGTGACTGGTGTTCAAAATTCAGGACATGCT-3’ and 5’-TTTCCAGTAGTTCCACTGACCACAAGTTTTA-3’, followed by verification of the mutation by sequencing.

### Three-Fragment Vector Construction

Plasmid DNA amplification and plasmid construction was performed with the MultiSite Gateway Three-Fragment Vector Construction kit (Invitrogen) using the primers 5’-GGGGACAACTTTGTATAG AAAAGTTGAGATCTCTAAAAGTTACATAAAATT-3’ and 5’-GGGGACTGCTTT TTTGTACAAACTTGCTGGAAAAGAAAATTTGATTTTTAA-3’ for amplifying the promoter region of *pie-1* gene. The amplified PCR product was cloned into pDONR P4-P1R using BP clonase creating the 5’ Gateway entry clone. The primers 5’-GGGGACAAGTTTGTACAAAAAAGCAGGCTACATGCCACCAGCAGTTCCAGCC-3’ and 5’-GGGGACCACTTTGTACAAGAAAGCTGGGTATTAGCGGACAGCCATGACAAT-3’ were used to amplify the *cct-4(vv39)* coding sequence from the plasmid made previously by *in-vitro* mutagenesis. The amplified gene was cloned into pDONR 221 to generate an internal Gateway entry clone. The primer pairs 5’-GGGGACAGCTTTCTTGTA CAAAGTGGAAATTTTCAGCATCTCGCGCCCGTG-3’ and 5’-GGGGACAACTTT GTATAATAAAGTTGGACTAGTAGGAAACAGTTATGTTTG-3’ were used to amplify the 3’ untranslated region of *unc-54.* The amplified product was cloned into pDONR P2R-P3 to create a 3’ Gateway entry clone. Combining all three entry clones into the Gateway compatible pCFJ150, a destination vector was constructed with the *pie-1* promoter, *cct-4(vv39)* gene, and *unc-54utr*.

### Mos Single Copy Insertion (MosSCI)

To generate a transgenic strain containing a single copy of *cct-4(vv39)*, MosSCI was performed as described in ^74^ using the ttT15605 MosSCI site insertion on chromosome II. 20-24h post L4 stage EG4322 (*ttTi5605* II*; unc-119(ed3)* III) adult nematodes were injected with pCFJ150 containing *pie-1p::cct-4(vv39)::unc-54utr*, together with two another constructs: pMR910 containing the GFP marker under the control of pharyngeal muscle promoter *myo-2* (*myo-2::GFP)*, and pJL44 containing Mos1 transposase under the control of the heat shock promoter (Phsp::transposase). The injected animals were individually plated and were left at 20°C overnight for recovery. On the next day the worms were heat shocked at 33-34°C for 1 hr, placed at 15°C for 4 hrs to recover, and moved to 20°C overnight. The worms were heat shocked again on the second and third day, and then were left at 20°C until starvation. Once starved, plates were screened for animals that showed rescue of the *unc-119(ed3)* phenotype and did not have the GFP marker. Transgenic animals were individually plated, and the presence of insertion was verified by PCR and sequencing.

### CRISPR-Cas9 Mutagenesis

CRISPR-mediated genome editing was performed as described in ^115^ using the N2 Bristol strain background, except for *htp-1(vv164)* which was made in *htp-2(tm2543)* null background. Injection mixes contained Cas9 protein (2 mg/ml; PNA Bio Inc), tracrRNA (8 µg/µl), *dpy-10* crRNA (8 µg/µl), *dpy-10* repair DNA template (500 ng/µl), gene of interest crRNA (8 µg/µl), repair DNA template (1 µg/µl), KCl (1M), Hepes pH 7.4 (200 mM). Young adults were injected and allowed to recover at 20°C. Individual Dpy or Roller F1 progeny were isolated on plates seeded with OP50 *E. coli* bacteria and screened for presence of mutation, confirmed by DNA sequencing were outcrossed three times. The HTP-3*^S37F^* (*htp-3[vv153]),* HTP-1*^S38F^ (htp-1[vv164] htp-2[tm2543]),* MDF-2*^S15F^ (mdf-2[vv154]), cct-4(vv151[ha::cct-4]), cct-5(vv169[flag::cct-5]),* and *cct-5(vv174[ha::cct-5])* genome edits were generated by CRISPR-Cas9 based site-directed mutagenesis supplemented with *dpy-10* co-conversion. The guide RNA and DNA template sequences used are listed in Supplemental Table 2.

### Preparation of protein extracts

Protein fractionation was performed as described in^96^ without modifications. Briefly, 10 confluent 100 mm Petri dishes containing synchronized young adults (24 hours post-L4) of the indicated genotypes were washed with M9 buffer and worms were collected by sedimentation. Worm pellet was washed extensively to remove residual bacteria, until a completely transparent supernatant was obtained. Excess M9 buffer was removed, and worms were resuspended in 4 mL of NP Buffer (10 mM HEPES-KOH pH 7.6, 1 mM EDTA, 10 mM KCl, 1.5 mM MgCl_2_, 0.25 mM sucrose, PMSF 1 mM and DTT 1 mM)/1 mL of worm pellet. The pellet was thoroughly mixed with NP buffer by placing tubes on a vortex for 10 seconds at maximum speed and then samples were immediately placed in liquid nitrogen. Tubes were stored at −80°C until lysis. After lysis and processing of the samples, cytosolic, nuclear soluble and chromatin bound fractions were produced: the chromatin bound fraction is obtained by incubation of nuclear pellet for 1 hour at 4°C with Benzonase (25U/100μl of nuclear extract). 30 μg of each fraction was employed for Western Blot analysis.

### Co-immunoprecipitation

Co-immunoprecipitation assays were performed as in^116^ without modifications. 1 mg of cytosolic extract was combined with 1 mg of nuclear extract (nuclear soluble and chromatin bound were pooled together) and incubated with pre-equilibrated agarose GFP traps (Chromotek, #gta-20) or magnetic FLAG traps (Sigma, #M8823) in buffer D (20 mM HEPES pH 7.9, 150 mM KCl, 20% glycerol, 0.2 mM EDTA, 0.2% Triton X-100) supplemented with complete protease inhibitor (Roche) overnight at 4°C. Beads were collected by centrifugation at 7500 rpm for 2 minutes (GFP traps) or with a magnetic rack (FLAG) at room temperature and extensively washed in buffer D. After the final wash, beads were resuspended in 2X Laemmli Buffer and boiled for 10 minutes. Eluted immunocomplexes were separated from the beads by centrifugation and loaded onto a precast 4%-20% gradient acrylamide gel (BioRad). After transfer onto a nitrocellulose membrane, blocking, primary and secondary antibodies incubation was carried out in 1X TBS containing 0.1% Tween and 5% milk for 1 hour at room temperature, overnight at 4°C and 2 hours at room temperature respectively. Antibodies used for western blot detection were: monoclonal mouse anti-HA (Cell Signaling, #2367) 1:1000, monoclonal rat anti-HA HRP-conjugated high affinity (Roche, #11867423001) 1:2000, monoclonal mouse anti-FLAG HRP-conjugated (Sigma, #A8592) 1:5000, polyclonal chicken anti-GFP (AbCam, #ab13970) 1:5000, monoclonal mouse anti-GAPDH (ThermoFisher, #AM4300) 1:5000, polyclonal rabbit anti Histone H3 (AbCam, #ab1791) 1:100,000. Western Blot detection was performed with Clarity Western ECL Substrate (BioRad, #1705060) and imaged on a G:Box chemidoc (Syngene).

## References

1. Zickler D, Kleckner N. Meiosis: Dances Between Homologs. Annu Rev Genet. 2023;57:1–63.

2. Moses MJ. Structure and function of the synaptonemal complex. Genetics. 1969;61(1):Suppl:41–51.

3. Hemmer LW, Blumenstiel JP. Holding it together: rapid evolution and positive selection in the synaptonemal complex of Drosophila. BMC Evol Biol. 2016;16:91.

4. Kursel LE, Cope HD, Rog O. Unconventional conservation reveals structure-function relationships in the synaptonemal complex. Elife. 2021;10.

5. Zickler D, Kleckner N. Meiotic chromosomes: integrating structure and function. Annu Rev Genet. 1999;33:603–754.

6. Hughes SE, Hawley RS. Alternative Synaptonemal Complex Structures: Too Much of a Good Thing? Trends Genet. 2020;36(11):833–44.

7. Rog O, Dernburg AF. Direct Visualization Reveals Kinetics of Meiotic Chromosome Synapsis. Cell Rep. 2015;10(10):1639–45.

8. Pollard MG, Rockmill B, Oke A, Anderson CM, Fung JC. Kinetic analysis of synaptonemal complex dynamics during meiosis of yeast Saccharomyces cerevisiae reveals biphasic growth and abortive disassembly. Front Cell Dev Biol. 2023;11:1098468.

9. Zickler D, Kleckner N. Recombination, Pairing, and Synapsis of Homologs during Meiosis. Cold Spring Harb Perspect Biol. 2015;7(6).

10. Phillips CM, Wong C, Bhalla N, Carlton PM, Weiser P, Meneely PM, et al. HIM-8 binds to the X chromosome pairing center and mediates chromosome-specific meiotic synapsis. Cell. 2005;123(6):1051–63.

11. Phillips CM, Dernburg AF. A family of zinc-finger proteins is required for chromosome-specific pairing and synapsis during meiosis in C. elegans. Dev Cell. 2006;11(6):817–29.

12. MacQueen AJ, Phillips CM, Bhalla N, Weiser P, Villeneuve AM, Dernburg AF. Chromosome sites play dual roles to establish homologous synapsis during meiosis in C. elegans. Cell. 2005;123(6):1037–50.

13. Sato A, Isaac B, Phillips CM, Rillo R, Carlton PM, Wynne DJ, et al. Cytoskeletal forces span the nuclear envelope to coordinate meiotic chromosome pairing and synapsis. Cell. 2009;139(5):907–19.

14. Penkner AM, Fridkin A, Gloggnitzer J, Baudrimont A, Machacek T, Woglar A, et al. Meiotic chromosome homology search involves modifications of the nuclear envelope protein Matefin/SUN-1. Cell. 2009;139(5):920–33.

15. Baudrimont A, Penkner A, Woglar A, Machacek T, Wegrostek C, Gloggnitzer J, et al. Leptotene/Zygotene Chromosome Movement Via the SUN/KASH Protein Bridge in. Plos Genet. 2010;6(11).

16. Wynne DJ, Rog O, Carlton PM, Dernburg AF. Dynein-dependent processive chromosome motions promote homologous pairing in C. elegans meiosis. J Cell Biol. 2012;196(1):47–64.

17. Labrador L, Barroso C, Lightfoot J, Muller-Reichert T, Flibotte S, Taylor J, et al. Chromosome movements promoted by the mitochondrial protein SPD-3 are required for homology search during Caenorhabditis elegans meiosis. Plos Genet. 2013;9(5):e1003497.

18. Olaya I, Burgess SM, Rog O. Formation and resolution of meiotic chromosome entanglements and interlocks. J Cell Sci. 2024;137(13).

19. Jitka Blazickova ST, Richard Bowman, Sowmya Sivakumar Geetha, Silma Subah, Sarit Smolikove, Verena Jantsch, Monique Zetka, Nicola Silva. Distinct intersecting pathways link homolog pairing to initiation of meiotic chromosome synapsis. bioRxiv 20240311584447. 2024.

20. Couteau F, Nabeshima K, Villeneuve A, Zetka M. A component of meiotic chromosome axes at the interface of homolog alignment, synapsis, nuclear reorganization, and recombination. Current Biology. 2004;14(7):585–92.

21. Couteau F, Zetka M. HTP-1 coordinates synaptonemal complex assembly with homolog alignment during meiosis in. Gene Dev. 2005;19(22):2744–56.

22. Martinez-Perez E, Villeneuve AM. HTP-1-dependent constraints coordinate homolog pairing and synapsis and promote chiasma formation during meiosis. Gene Dev. 2005;19(22):2727–43.

23. Goodyer W, Kaitna S, Couteau F, Ward JD, Boulton SJ, Zetka M. HTP-3 links DSB formation with homolog pairing and crossing over during meiosis. Dev Cell. 2008;14(2):263–74.

24. Bohr T, Ashley G, Eggleston E, Firestone K, Bhalla N. Synaptonemal Complex Components Are Required for Meiotic Checkpoint Function in Caenorhabditis elegans. Genetics. 2016;204(3):987–97.

25. Grey C, de Massy B. Chromosome Organization in Early Meiotic Prophase. Front Cell Dev Biol. 2021;9:688878.

26. Ur SN, Corbett KD. Architecture and Dynamics of Meiotic Chromosomes. Annu Rev Genet. 2021;55:497–526.

27. Prince JP, Martinez-Perez E. Functions and Regulation of Meiotic HORMA-Domain Proteins. Genes (Basel). 2022;13(5).

28. Aravind L, Koonin EV. The HORMA domain: a common structural denominator in mitotic checkpoints, chromosome synapsis and DNA repair. Trends Biochem Sci. 1998;23(8):284–6.

29. Rosenberg SC, Corbett KD. The multifaceted roles of the HORMA domain in cellular signaling. J Cell Biol. 2015;211(4):745–55.

30. Luo X, Tang Z, Xia G, Wassmann K, Matsumoto T, Rizo J, et al. The Mad2 spindle checkpoint protein has two distinct natively folded states. Nat Struct Mol Biol. 2004;11(4):338–45.

31. Gu Y, Desai A, Corbett KD. Evolutionary Dynamics and Molecular Mechanisms of HORMA Domain Protein Signaling. Annu Rev Biochem. 2022;91:541–69.

32. West AMV, Komives EA, Corbett KD. Conformational dynamics of the Hop1 HORMA domain reveal a common mechanism with the spindle checkpoint protein Mad2. Nucleic Acids Res. 2018;46(1):279–92.

33. Hollingsworth NM, Byers B. HOP1: a yeast meiotic pairing gene. Genetics. 1989;121(3):445–62.

34. Schwacha A, Kleckner N. Identification of joint molecules that form frequently between homologs but rarely between sister chromatids during yeast meiosis. Cell. 1994;76(1):51–63.

35. Woltering D, Baumgartner B, Bagchi S, Larkin B, Loidl J, de los Santos T, et al. Meiotic segregation, synapsis, and recombination checkpoint functions require physical interaction between the chromosomal proteins Red1p and Hop1p. Mol Cell Biol. 2000;20(18):6646–58.

36. Niu H, Wan L, Baumgartner B, Schaefer D, Loidl J, Hollingsworth NM. Partner choice during meiosis is regulated by Hop1-promoted dimerization of Mek1. Mol Biol Cell. 2005;16(12):5804–18.

37. Shin YH, Choi Y, Erdin SU, Yatsenko SA, Kloc M, Yang F, et al. Hormad1 mutation disrupts synaptonemal complex formation, recombination, and chromosome segregation in mammalian meiosis. Plos Genet. 2010;6(11):e1001190.

38. Daniel K, Lange J, Hached K, Fu J, Anastassiadis K, Roig I, et al. Meiotic homologue alignment and its quality surveillance are controlled by mouse HORMAD1. Nat Cell Biol. 2011;13(5):599–610.

39. Kogo H, Tsutsumi M, Inagaki H, Ohye T, Kiyonari H, Kurahashi H. HORMAD2 is essential for synapsis surveillance during meiotic prophase via the recruitment of ATR activity. Genes Cells. 2012;17(11):897–912.

40. Wojtasz L, Cloutier JM, Baumann M, Daniel K, Varga J, Fu J, et al. Meiotic DNA double-strand breaks and chromosome asynapsis in mice are monitored by distinct HORMAD2-independent and -dependent mechanisms. Genes Dev. 2012;26(9):958–73.

41. Sanchez-Moran E, Santos JL, Jones GH, Franklin FC. ASY1 mediates AtDMC1-dependent interhomolog recombination during meiosis in Arabidopsis. Genes Dev. 2007;21(17):2220–33.

42. Kim Y, Rosenberg SC, Kugel CL, Kostow N, Rog O, Davydov V, et al. The chromosome axis controls meiotic events through a hierarchical assembly of HORMA domain proteins. Dev Cell. 2014;31(4):487–502.

43. Kim Y, Kostow N, Dernburg AF. The Chromosome Axis Mediates Feedback Control of CHK-2 to Ensure Crossover Formation in C. elegans. Dev Cell. 2015;35(2):247–61.

44. Ferrandiz N, Barroso C, Telecan O, Shao N, Kim HM, Testori S, et al. Spatiotemporal regulation of Aurora B recruitment ensures release of cohesion during C. elegans oocyte meiosis. Nat Commun. 2018;9(1):834.

45. Martinez-Perez E, Schvarzstein M, Barroso C, Lightfoot J, Dernburg AF, Villeneuve AM. Crossovers trigger a remodeling of meiotic chromosome axis composition that is linked to two-step loss of sister chromatid cohesion. Gene Dev. 2008;22(20):2886–901.

46. Severson AF, Ling L, van Zuylen V, Meyer BJ. The axial element protein HTP-3 promotes cohesin loading and meiotic axis assembly in C. elegans to implement the meiotic program of chromosome segregation. Genes Dev. 2009;23(15):1763–78.

47. Goodyer W, Kaitna S, Couteau F, Ward JD, Boulton SJ, Zetka M. HTP-3 links DSB formation with homolog pairing and crossing over during C. elegans meiosis. Dev Cell. 2008;14(2):263–74.

48. Zetka MC, Kawasaki I, Strome S, Müller F. Synapsis acid chiasma formation in require HIM-3, a meiotic chromosome core component that functions in chromosome segregation. Gene Dev. 1999;13(17):2258–70.

49. Couteau F, Nabeshima K, Villeneuve A, Zetka M. A component of C. elegans meiotic chromosome axes at the interface of homolog alignment, synapsis, nuclear reorganization, and recombination. Curr Biol. 2004;14(7):585–92.

50. Consuelo Barroso JPP, Punam Rattu, Daimona Kundé, Nuria Ferrandiz, Syma Khalid, Enrique Martinez-Perez. Two structurally mobile regions control the conformation and function of metamorphic meiotic HORMAD proteins. bioRxiv 20240805606648. 2024.

51. Vader G. Pch2(TRIP13): controlling cell division through regulation of HORMA domains. Chromosoma. 2015;124(3):333–9.

52. Bhalla N. PCH-2 and meiotic HORMADs: A module for evolutionary innovation in meiosis? Curr Top Dev Biol. 2023;151:317–44.

53. Joshi N, Barot A, Jamison C, Borner GV. Pch2 links chromosome axis remodeling at future crossover sites and crossover distribution during yeast meiosis. Plos Genet. 2009;5(7):e1000557.

54. Miao C, Tang D, Zhang H, Wang M, Li Y, Tang S, et al. Central region component1, a novel synaptonemal complex component, is essential for meiotic recombination initiation in rice. Plant Cell. 2013;25(8):2998–3009.

55. Lambing C, Osman K, Nuntasoontorn K, West A, Higgins JD, Copenhaver GP, et al. Arabidopsis PCH2 Mediates Meiotic Chromosome Remodeling and Maturation of Crossovers. Plos Genet. 2015;11(7):e1005372.

56. Chotiner JY, Leu NA, Yang F, Cossu IG, Guan Y, Lin H, et al. TRIP13 localizes to synapsed chromosomes and functions as a dosage-sensitive regulator of meiosis. Elife. 2024;12.

57. Deshong AJ, Ye AL, Lamelza P, Bhalla N. A quality control mechanism coordinates meiotic prophase events to promote crossover assurance. Plos Genet. 2014;10(4):e1004291.

58. Lopez T, Dalton K, Frydman J. The Mechanism and Function of Group II Chaperonins. J Mol Biol. 2015;427(18):2919–30.

59. Yam AY, Xia Y, Lin HT, Burlingame A, Gerstein M, Frydman J. Defining the TRiC/CCT interactome links chaperonin function to stabilization of newly made proteins with complex topologies. Nat Struct Mol Biol. 2008;15(12):1255–62.

60. Russmann F, Stemp MJ, Monkemeyer L, Etchells SA, Bracher A, Hartl FU. Folding of large multidomain proteins by partial encapsulation in the chaperonin TRiC/CCT. Proc Natl Acad Sci U S A. 2012;109(52):21208–15.

61. Zeng C, Han S, Pan Y, Huang Z, Zhang B, Zhang B. Revisiting the chaperonin T-complex protein-1 ring complex in human health and disease: A proteostasis modulator and beyond. Clin Transl Med. 2024;14(2):e1592.

62. Huang R, Yu M, Li CY, Zhan YQ, Xu WX, Xu F, et al. New insights into the functions and localization of nuclear CCT protein complex in K562 leukemia cells. Proteomics Clin Appl. 2012;6(9-10):467–75.

63. Joly EC, Tremblay E, Tanguay RM, Wu Y, Bibor-Hardy V. TRiC-P5, a novel TCP1-related protein, is localized in the cytoplasm and in the nuclear matrix. J Cell Sci. 1994;107 (Pt 10):2851–9.

64. Gerner C, Holzmann K, Meissner M, Gotzmann J, Grimm R, Sauermann G. Reassembling proteins and chaperones in human nuclear matrix protein fractions. J Cell Biochem. 1999;74(2):145–51.

65. Roobol A, Carden MJ. Subunits of the eukaryotic cytosolic chaperonin CCT do not always behave as components of a uniform hetero-oligomeric particle. Eur J Cell Biol. 1999;78(1):21–32.

66. Gerner C, Gotzmann J, Frohwein U, Schamberger C, Ellinger A, Sauermann G. Proteome analysis of nuclear matrix proteins during apoptotic chromatin condensation. Cell Death Differ. 2002;9(6):671–81.

67. Kasembeli M, Lau WC, Roh SH, Eckols TK, Frydman J, Chiu W, et al. Modulation of STAT3 folding and function by TRiC/CCT chaperonin. PLoS Biol. 2014;12(4):e1001844.

68. Saegusa K, Sato M, Sato K, Nakajima-Shimada J, Harada A, Sato K. Caenorhabditis elegans chaperonin CCT/TRiC is required for actin and tubulin biogenesis and microvillus formation in intestinal epithelial cells. Mol Biol Cell. 2014;25(20):3095–104.

69. Soues S, Kann ML, Fouquet JP, Melki R. The cytosolic chaperonin CCT associates to cytoplasmic microtubular structures during mammalian spermiogenesis and to heterochromatin in germline and somatic cells. Exp Cell Res. 2003;288(2):363–73.

70. Gvozdenov Z, Kolhe J, Freeman BC. The Nuclear and DNA-Associated Molecular Chaperone Network. Cold Spring Harb Perspect Biol. 2019;11(10).

71. Zlata Gvozdenov AYTP, Anusmita Biswas, Zeno Barcutean, Daniel Gestaut, Judith Frydman, Kevin Struhl, Brian C Freeman. TRiC/CCT Chaperonin Governs RNA Polymerase II Activity in the Nucleus to Support RNA Homeostasis. 2024.

72. Dekker C, Stirling PC, McCormack EA, Filmore H, Paul A, Brost RL, et al. The interaction network of the chaperonin CCT. EMBO J. 2008;27(13):1827–39.

73. Wicks SR, Yeh RT, Gish WR, Waterston RH, Plasterk RH. Rapid gene mapping in Caenorhabditis elegans using a high density polymorphism map. Nat Genet. 2001;28(2):160–4.

74. Frokjaer-Jensen C, Davis MW, Hopkins CE, Newman BJ, Thummel JM, Olesen SP, et al. Single-copy insertion of transgenes in Caenorhabditis elegans. Nat Genet. 2008;40(11):1375–83.

75. Cong Y, Baker ML, Jakana J, Woolford D, Miller EJ, Reissmann S, et al. 4.0-A resolution cryo-EM structure of the mammalian chaperonin TRiC/CCT reveals its unique subunit arrangement. Proc Natl Acad Sci U S A. 2010;107(11):4967–72.

76. Dernburg AF, McDonald K, Moulder G, Barstead R, Dresser M, Villeneuve AM. Meiotic recombination initiates by a conserved mechanism and is dispensable for homologous chromosome synapsis. Cell. 1998;94(3):387–98.

77. MacQueen AJ, Colaiacovo MP, McDonald K, Villeneuve AM. Synapsis-dependent and - independent mechanisms stabilize homolog pairing during meiotic prophase in C. elegans. Genes Dev. 2002;16(18):2428–42.

78. Zetka M, Paouneskou D, Jantsch V. “The nuclear envelope, a meiotic jack-of-all-trades”. Curr Opin Cell Biol. 2020;64:34–42.

79. Harper NC, Rillo R, Jover-Gil S, Assaf ZJ, Bhalla N, Dernburg AF. Pairing centers recruit a Polo-like kinase to orchestrate meiotic chromosome dynamics in C. elegans. Dev Cell. 2011;21(5):934–47.

80. Labella S, Woglar A, Jantsch V, Zetka M. Polo Kinases Establish Links between Meiotic Chromosomes and Cytoskeletal Forces Essential for Homo log Pairing. Dev Cell. 2011;21(5):948–58.

81. MacQueen AJ, Villeneuve AM. Nuclear reorganization and homologous chromosome pairing during meiotic prophase require C. elegans chk-2. Genes Dev. 2001;15(13):1674–87.

82. Joachimiak LA, Walzthoeni T, Liu CW, Aebersold R, Frydman J. The structural basis of substrate recognition by the eukaryotic chaperonin TRiC/CCT. Cell. 2014;159(5):1042–55.

83. Kelly JJ, Tranter D, Pardon E, Chi G, Kramer H, Happonen L, et al. Snapshots of actin and tubulin folding inside the TRiC chaperonin. Nat Struct Mol Biol. 2022;29(5):420–9.

84. Almutairi ZM. Comparative genomics of HORMA domain-containing proteins in prokaryotes and eukaryotes. Cell Cycle. 2018;17(23):2531–46.

85. Cuellar J, Ludlam WG, Tensmeyer NC, Aoba T, Dhavale M, Santiago C, et al. Structural and functional analysis of the role of the chaperonin CCT in mTOR complex assembly. Nat Commun. 2019;10(1):2865.

86. Gao YJ, Thomas JO, Chow RL, Lee GH, Cowan NJ. A Cytoplasmic Chaperonin That Catalyzes Beta-Actin Folding. Cell. 1992;69(6):1043–50.

87. Yaffe MB, Farr GW, Miklos D, Horwich AL, Sternlicht ML, Sternlicht H. TCP1 complex is a molecular chaperone in tubulin biogenesis. Nature. 1992;358(6383):245–8.

88. Lundin VF, Srayko M, Hyman AA, Leroux MR. Efficient chaperone-mediated tubulin biogenesis is essential for cell division and cell migration in C. elegans. Dev Biol. 2008;313(1):320–34.

89. Camasses A, Bogdanova A, Shevchenko A, Zachariae W. The CCT chaperonin promotes activation of the anaphase-promoting complex through the generation of functional Cdc20. Mol Cell. 2003;12(1):87–100.

90. Hubstenberger A, Cameron C, Noble SL, Keenan S, Evans TC. Modifiers of solid RNP granules control normal RNP dynamics and mRNA activity in early development. J Cell Biol. 2015;211(3):703–16.

91. Elaswad MT, Gao M, Tice VE, Bright CG, Thomas GM, Munderloh C, et al. The CCT chaperonin and actin modulate the ER and RNA-binding protein condensation during oogenesis and maintain translational repression of maternal mRNA and oocyte quality. Mol Biol Cell. 2024;35(10):ar131.

92. Ma X, Lu C, Chen Y, Li S, Ma N, Tao X, et al. CCT2 is an aggrephagy receptor for clearance of solid protein aggregates. Cell. 2022;185(8):1325–45 e22.

93. Leroux MR, Candido EP. Molecular analysis of Caenorhabditis elegans tcp-1, a gene encoding a chaperonin protein. Gene. 1995;156(2):241–6.

94. Leroux MR, Ma BJ, Batelier G, Melki R, Candido EP. Unique structural features of a novel class of small heat shock proteins. J Biol Chem. 1997;272(19):12847–53.

95. Noormohammadi A, Khodakarami A, Gutierrez-Garcia R, Lee HJ, Koyuncu S, Konig T, et al. Somatic increase of CCT8 mimics proteostasis of human pluripotent stem cells and extends C. elegans lifespan. Nat Commun. 2016;7:13649.

96. Silva N, Ferrandiz N, Barroso C, Tognetti S, Lightfoot J, Telecan O, et al. The Fidelity of Synaptonemal Complex Assembly Is Regulated by a Signaling Mechanism that Controls Early Meiotic Progression. Dev Cell. 2014;31(4):503–11.

97. Guisbert E, Czyz DM, Richter K, McMullen PD, Morimoto RI. Identification of a tissue-selective heat shock response regulatory network. Plos Genet. 2013;9(4):e1003466.

98. Maheshwari R, Rahman MM, Joseph-Strauss D, Cohen-Fix O. An RNAi screen for genes that affect nuclear morphology in Caenorhabditis elegans reveals the involvement of unexpected processes. G3 (Bethesda). 2021;11(11).

99. Chen Y, Kang J, Zhen R, Zhang L, Chen C. A genome-wide CRISPR screen identifies the CCT chaperonin as a critical regulator of vesicle trafficking. FASEB J. 2023;37(2):e22757.

100. Ahuja JS, Sandhu R, Mainpal R, Lawson C, Henley H, Hunt PA, et al. Control of meiotic pairing and recombination by chromosomally tethered 26S proteasome. Science. 2017;355(6323):408–11.

101. Gestaut D, Zhao Y, Park J, Ma B, Leitner A, Collier M, et al. Structural visualization of the tubulin folding pathway directed by human chaperonin TRiC/CCT. Cell. 2022;185(25):4770–87 e20.

102. Rog O, Kohler S, Dernburg AF. The synaptonemal complex has liquid crystalline properties and spatially regulates meiotic recombination factors. Elife. 2017;6.

103. Castellano-Pozo M, Pacheco S, Sioutas G, Jaso-Tamame AL, Dore MH, Karimi MM, et al. Surveillance of cohesin-supported chromosome structure controls meiotic progression. Nat Commun. 2020;11(1):4345.

104. Yang C, Hu B, Portheine SM, Chuenban P, Schnittger A. State changes of the HORMA protein ASY1 are mediated by an interplay between its closure motif and PCH2. Nucleic Acids Res. 2020;48(20):11521–35.

105. Herruzo E, Lago-Maciel A, Baztan S, Santos B, Carballo JA, San-Segundo PA. Pch2 orchestrates the meiotic recombination checkpoint from the cytoplasm. Plos Genet. 2021;17(7):e1009560.

106. Banks CAS, Miah S, Adams MK, Eubanks CG, Thornton JL, Florens L, et al. Differential HDAC1/2 network analysis reveals a role for prefoldin/CCT in HDAC1/2 complex assembly. Sci Rep. 2018;8(1):13712.

107. Dekker C. On the role of the chaperonin CCT in the just-in-time assembly process of APC/CCdc20. Febs Lett. 2010;584(3):477–81.

108. Rao HB, Qiao H, Bhatt SK, Bailey LR, Tran HD, Bourne SL, et al. A SUMO-ubiquitin relay recruits proteasomes to chromosome axes to regulate meiotic recombination. Science. 2017;355(6323):403-7.

109. Brenner S. The genetics of Caenorhabditis elegans. Genetics. 1974;77(1):71–94.

110. Colaiacovo MP, MacQueen AJ, Martinez-Perez E, McDonald K, Adamo A, La Volpe A, et al. Synaptonemal complex assembly in C. elegans is dispensable for loading strand-exchange proteins but critical for proper completion of recombination. Dev Cell. 2003;5(3):463–74.

111. Devigne AP, Bhalla N. Mad1’s ability to interact with Mad2 is essential to regulate and monitor meiotic synapsis in C. elegans. Plos Genet. 2021;17(11).

112. Zalevsky J, MacQueen AJ, Duffy JB, Kemphues KJ, Villeneuve AM. Crossing over during Caenorhabditis elegans meiosis requires a conserved MutS-based pathway that is partially dispensable in budding yeast. Genetics. 1999;153(3):1271–83.

113. Schindelin J, Arganda-Carreras I, Frise E, Kaynig V, Longair M, Pietzsch T, et al. Fiji: an open-source platform for biological-image analysis. Nat Methods. 2012;9(7):676–82.

114. Fire A, Xu S, Montgomery MK, Kostas SA, Driver SE, Mello CC. Potent and specific genetic interference by double-stranded RNA in Caenorhabditis elegans. Nature. 1998;391(6669):806–11.

115. Paix A, Folkmann A, Seydoux G. Precision genome editing using CRISPR-Cas9 and linear repair templates in C. elegans. Methods. 2017;121-122:86–93.

116. Janisiw E, Dello Stritto MR, Jantsch V, Silva N. BRCA1-BARD1 associate with the synaptonemal complex and pro-crossover factors and influence RAD-51 dynamics during Caenorhabditis elegans meiosis. Plos Genet. 2018;14(11):e1007653.

117. Tunyasuvunakool K, Adler J, Wu Z, Green T, Zielinski M, Zidek A, et al. Highly accurate protein structure prediction for the human proteome. Nature. 2021;596(7873):590–6.

