## Supplementary material for "A nuclear TRiC/CCT chaperonin assembles meiotic HORMAD proteins into chromosome axes competent for crossing over": Supplemetal Data

**Supplemental Figure Legends**

**Figure S1. Conservation of P382 in TRiC subunits**. M-COFFEE alignments of *C. elegans* CCT-4 and **A)** the CCT4 subunit of the indicated species and **B)** with the other seven TRiC subunits in *C. elegans*; proline is only present in CCT-4 and not in other subunits.

**Figure S2. *cct-4(RNAi)* disrupts germ cell proliferation and formation of the transition zone.** Immunofluorescence micrographs showing DAPI-stained hermaphrodite germ lines. **A)** *rrf-1(pk1417)* mutants show wild-type germline organization and progression through meiotic prophase. **B)** *rrf-1(pk1417); cct-4(RNAi)* germlines are smaller with fewer germ cells (distal end is at bottom) and are marked by the appearance of micronuclei (inset, orange arrows) that are hallmarks of defects in mitotic processes. A transition zone is defined by the presence of nuclei with the chromatin polarization characteristic of leptotene/zygotene is not present in *rrf-1(pk1417); cct-4(RNAi)* germlines. Scale bars, 5μm.

**Figure S3. α-CCT-4 antibody specificity.** DNA, HTP-3, and CCT-4 staining of nuclei from the pachytene region of wild type and *cct-4(RNAi)* germlines*.* In *cct-4(RNAi)* germlines, CCT-4 is not detectable in the nuclei and is severely reduced in cytoplasm*.* Scale bars, 5μm.

**Figure S4. CCT-5 form a complex with CCT-4 and HIM-3 *in vivo*. A)** Western blot analysis showing that FLAG::CCT-5 is abundantly pulled down by HA::CCT-4 immunoprecipitation. **B)** FLAG::CCT-5 establishes a physical interaction with GFP::HIM-3. **C)** Western blot analysis with the indicated antibodies on protein extracts used as the inputs for the above-mentioned co-immunoprecipitation experiments. Equal amounts of protein extracts were loaded.

**Table S1. Strains used in this study.**

| STRAIN | GENOTYPE |
|---|---|
| N2 | Wild type var. Bristol |
| ATG-1 | *htp-1(gk174) htp-2(tm2543) IV* |
| AV271 | *him-3(me80) IV* |
| CA1282 | *him-3[ie114(gfp::him-3)] IV* |
| CB138 | *unc-24(e138) II* |
| CB1147 | *him-3(e1147) IV* |
| CB1256 | *him-3(e1256) IV* |
| CB4856 | Wild type var. Hawaiian |
| EZ35 | *him-3(vv6) IV* |
| EZ68 | *him-3(gk149) IV* |
| EZ73 | *him-3(vv6) unc-24(e138) IV* |
| EZ88 | *him-3(vv6) unc-24(e138) IV; cct-4(vv39) II* |
| EZ143 | *cct-4(vv39) II* |
| EZ150 | *him-3(vv6) IV; cct-4(vv39) II* |
| EZ302 | *unc-119(ed3) III; him-3(vv6) IV; ojls7[zyg-12A::GFP + unc-119(+)]?* |
| EZ501 | *htp-3(vv153) I* |
| EZ507 | *htp-3(vv153) I; cct-4(vv39) II* |
| EZ529 | *htp-1(vv164) htp-2(tm2543) IV* |
| EZ533 | *cct-4(vv151[ha::cct-4]) II* |
| EZ562 | *cct-5(vv169[flag::cct-5]) III* |
| EZ563 | *cct-5(vv169(flag::cct-5]) III; him-3(ie114(gfp::him-3)] IV* |
| EZ576 | *htp-1(vv164) htp-2(tm2543) IV; cct-4(vv39) II* |
| EZ599 | *cct-4(vv151[HA:cct-4]) II; gfp::him-3 IV* |
| FX30203 | *tmC25(unc-5[tmIs1241]) IV* |
| TY5038 | *htp-3(tm3655)I / hT2 [bli-4(e937) let-? (q782) qIs48] I;III* |
| WH220 | *ojIs7[zyg-12A::GFP + unc-119(+)]?* |

**Table S2. CRISPR materials for mutant alleles made in this study.**

| Allele | Guide RNA and DNA repair templates | Genotyping primers and detection RE |
|---|---|---|
| *htp-3(vv153)* | 5' GTT TGC AAG ATA AAC GCA GT 3'<br><br>5'TTCAGGCTATATTCAACGAGCAAA CATTGAAAAATGGTGATGAAAATTC GAAAAGCTTTCTTGAAGTGATGGCT AACTGCGTTTATCTTGCAAACTCAA CAATTCTCCGTGAACG 3' | Forward primer: CTT CCA TGG GTC TCG CCG<br>Reverse primer: GAC ACA ACT CCG TCC TTC TGG<br><br>HaeIII |
| *htp-1(vv164)* | 5' AAA CGC GAC GTA GAT TGC TC 3'<br><br>5'CATCTTTAGAAATGGTCCAAACTTT TCCCAAGGATTGTCAGCGACCCGGA TCGATTCTCCAATTTCATGACTAGAG CAATCTACGTCGCGTTTTCGGCTGTT CTTAGGAACCGTA 3' | Forward primer: CGC CCT TGG AAA CCA TCT AC<br>Reverse primer: GTC ATG CAT TTC AGC TTT TCT GA<br><br>HinfI |
| *cct-4[vv151(ha::cct-4)]* | 5' GCT GCG GCT GGA ACT GCT GG 3'<br><br>5'TCAACATCATAATTTAATTTGCAG ATGTACCCATATGATGTCCCGGATT ACGCTTACCCATATGATGTCCCGGA TTACGCTCCACCAGCAGTTCCAGCC GCAGCTGCAACAGCCCGACAATCG GCTTCCGGTCGCGAGCGCAATTTCA AGGATAAGG 3' | Forward primer: TCG CCA CGA AAC CAG TATTT<br>Reverse primer: TTC GAC AAG CTT CTT CAG<br><br>PfoI |
| *cct-5[vv169(flag::cct-5)]* | 5' GTG TCT GTC AAG ACG GAA CT 3'<br><br>5'CTAATATTTTCAGTGTCTGTCAAGA CGGAACTTGGATAAGAAAATGGACT ACAAGGACGACGACGACAAGGCGCA GTCATCTGCGCAACTACTGTTCGATG AGAGCGGGCAACCG 3' | Forward primer: GCG GGT TTG ATA TCG AGA ACA TT<br>Reverse primer: CCG GTG ATA CGC TTC TGG TT |
| *dpy-10* | 5' GCU ACC AUA GGC ACC ACG AGG UUU UAG AGC UAU GCU 3'<br><br>5' CACTTGAACTTCAATACGGCAAGA TGAGAATGACTGGAAACCGTACCGC ATGCGGTGCCTATGGTAGCGGAGC TTCACATGGCTTCAGACCAACAGCCT AT 3' | N/A |

Figure S1

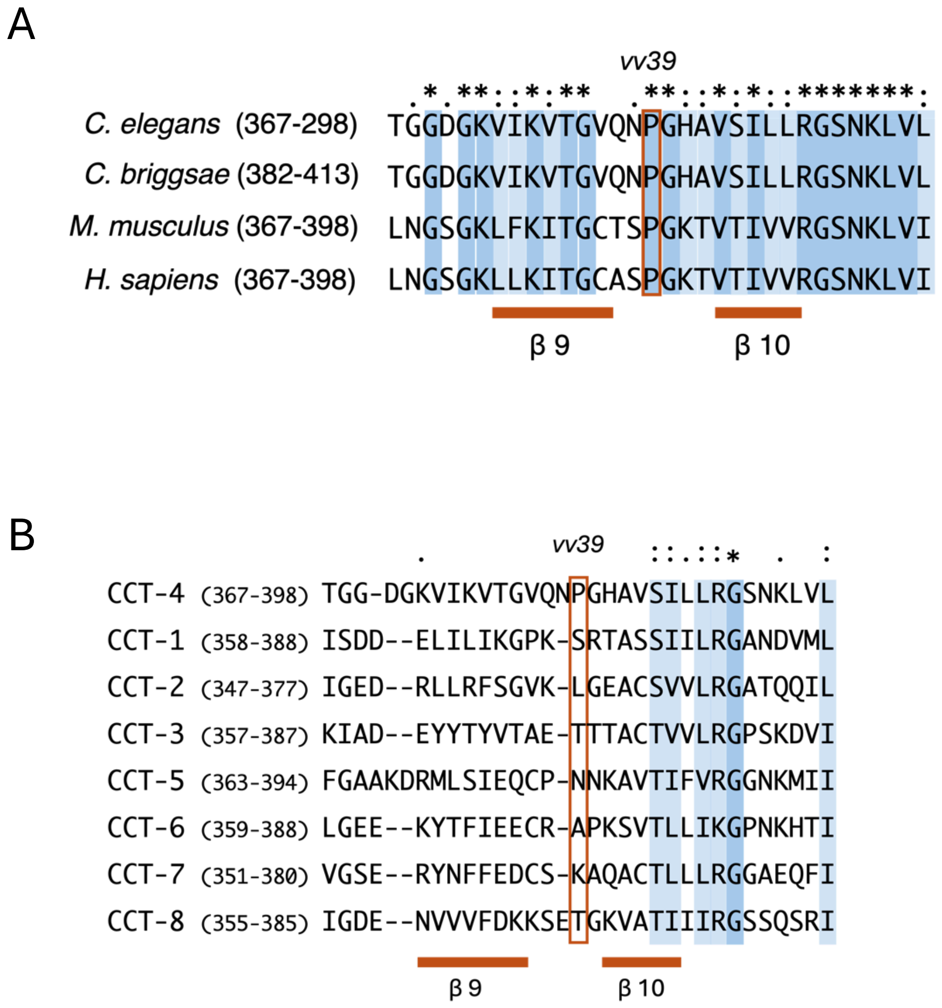
A
vv39
C. elegans (367-298) TGGDGKVIKVTGVQNPGHAVSILLRGSNKLVL
C. briggsae (382-413) TGGDGKVIKVTGVQNPGHAVSILLRGSNKLVL
M. musculus (367-398) LNGSGKLFKITGCTSPGKTVTIVVRGSNKLVI
H. sapiens (367-398) LNGSGKLLKITGCASPGKTVTIVVRGSNKLVI
β 9
β 10
B
vv39
CCT-4 (367-398) TGG-DGKVIKVTGVQNPGHAVSILLRGSNKLVL
CCT-1 (358-388) ISDD--ELILIKGPK-SRTASSIILRGANDVML
CCT-2 (347-377) IGED--RLLRFSGVK-LGEACSVVLRGATQQIL
CCT-3 (357-387) KIAD--EYYTYVTAE-TTTACTVVLRGPSKDVI
CCT-5 (363-394) FGAAKDRMLSIEQCP-NNKAVTIFVRGGNKMII
CCT-6 (359-388) LGEE--KYTFIEECR-APKSVTLLIKGPNKHTI
CCT-7 (351-380) VGSE--RYNFFEDCS-KAQACTLLLRGGAEQFI
CCT-8 (355-385) IGDE--NVVVFDKKSETGKVATIIIRGSSQSRI
β 9
β 10

Figure S2

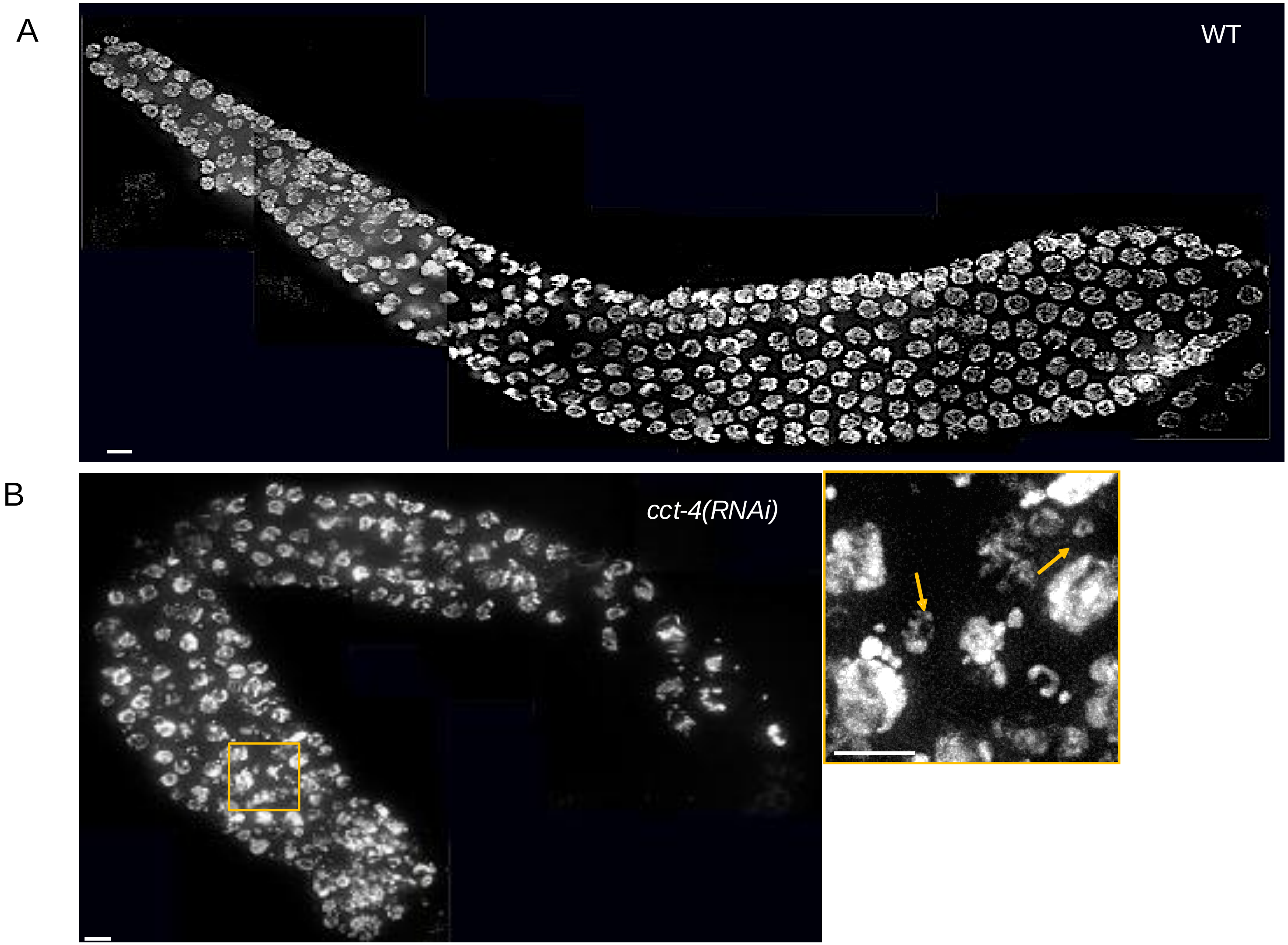

Figure S3

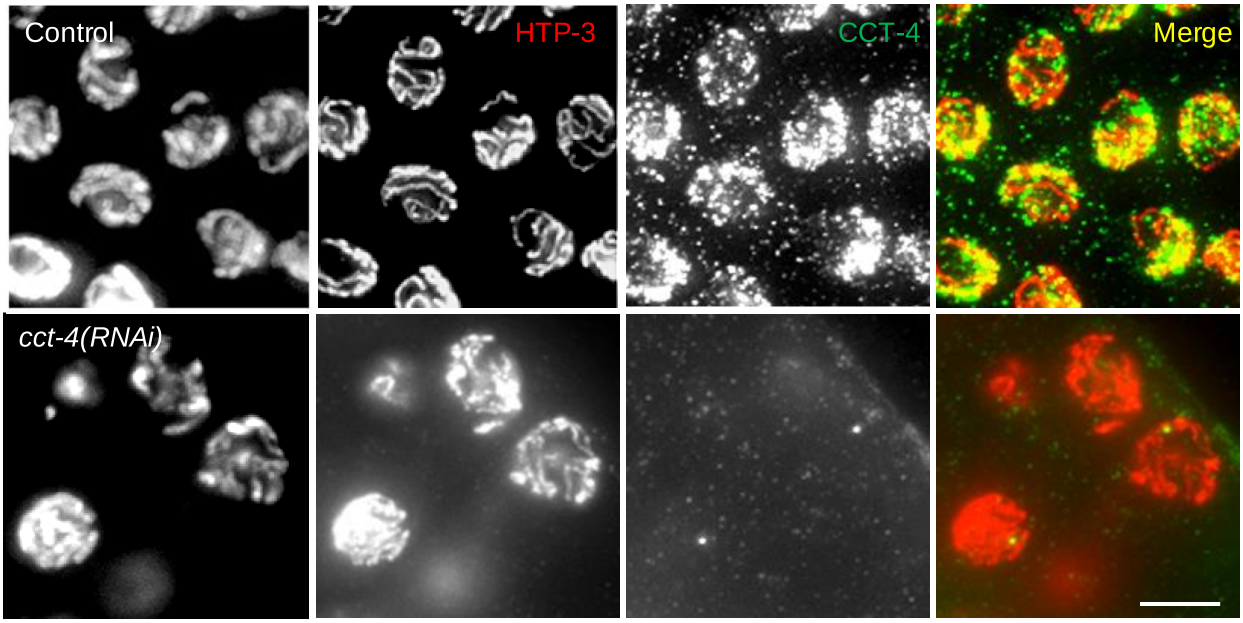

Figure S4

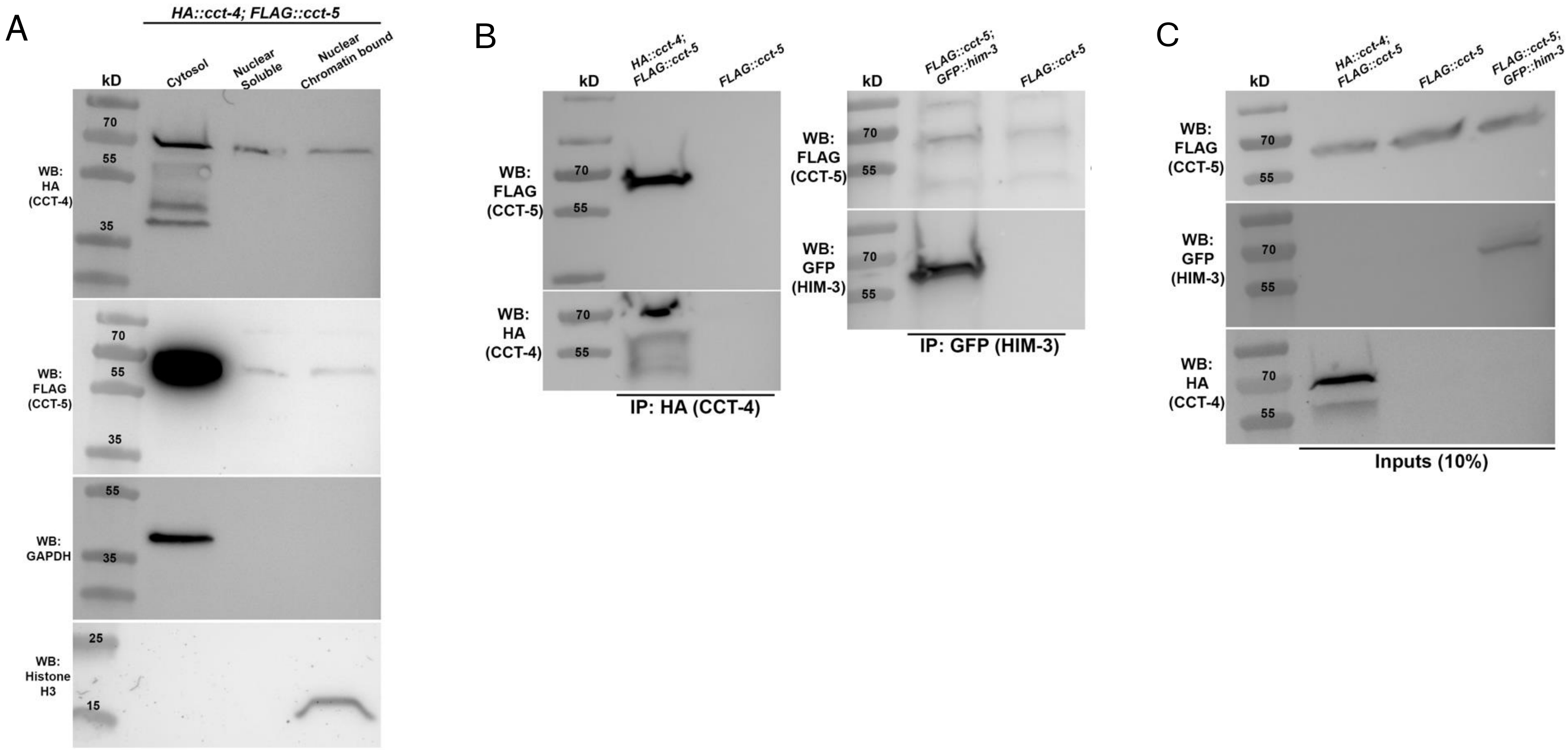
A
HA::cct-4; FLAG::cct-5
kD
Cytosol
Nuclear Soluble
Nuclear Chromatin bound
70
55
35
25
15
WB: HA (CCT-4)
WB: FLAG (CCT-5)
WB: GAPDH
WB: Histone H3
B
HA::cct-4; FLAG::cct-5
FLAG::cct-5
FLAG::cct-5; GFP::him-3
WB: FLAG (CCT-5)
WB: HA (CCT-4)
WB: GFP (HIM-3)
IP: HA (CCT-4)
IP: GFP (HIM-3)
C
Inputs (10%)
